# 4D confocal microscopy of time dependent cell and nuclear osmotic response reveals differential osmoregulation and transport kinetics

**DOI:** 10.64898/2026.09.30.755591

**Authors:** Iqra Azam, Dominic J. Olver, James D. Benson

**Author notes:** Corresponding author: James D. Benson.

## Abstract

A cell’s ability to maintain its volume is a prerequisite for its essential physiological functions. Changes in extracellular osmolarity induce the flux of water and solutes across the cell and nuclear membrane and, therefore, alters cell and nuclear volumes. Here, we investigated the equilibrium and dynamic volumetric responses of cells and nuclei in adherent human hepatoma (HepG2) cell monolayers using live three-dimensional imaging. Through image segmentation, 3D reconstruction, and volume rendering, precise cell and nuclear volumes were quantified. We found that during hypoosmotic stress, cells swell and lose osmolytes through active volume regulation, and nuclei undergo greater relative expansion than cells. During hyperosmotic stress, we found that both cells and nuclei shrink rapidly, develop irregular morphologies and undergo passive volume regulation. Furthermore, we explored the osmotic responses in these cells by restoring isosmotic conditions following osmotic stress and found that nuclei returned to volumes greater than their initial isosmotic volume, whereas cytoplasmic volume returned to a value below its initial isosmotic volume following swelling. By fitting coupled transport models water and solute permeability estimates for both cells and nuclei were made from their dynamic responses to different anisosmotic media with and without the presence of permeating solutes dimethyl sulfoxide and ethylene glycol. Together, these quantitative measurements of cell, cytoplasm, and nuclear volumetric responses and associated transport parameters have important implications for understanding cell physiology and the mechanism of osmoregulation.

## 1. Introduction

Accurate estimation of physicochemical parameters and cell volume is important for understanding the biological responses of cells to different mechanical or osmotic stresses [1, 2]. It has been demonstrated that a variety of cell types show physiological responses to physical stimuli, including mechanical deformation [3], hydrostatic pressure, fluid shear stress, and osmotic stress [4] [5]. In response to these stresses, ion channels and other membrane-bound receptors may be activated, altering cell behaviour [6]. Furthermore, the nucleus and other intracellular organelles can also be affected by physical stress at the subcellular level as mechanical or osmotic stress propagates through the cell [7]. However, how extracellular osmotic stress differentially affects the cell and nucleus, and how the inherent physicochemical properties of nucleus influence these responses remains poorly understood. A better understanding of the biophysical properties of the cells and nuclei may therefore help elucidate the significance of these stress-induced physical changes.

A standard assumption, known as the Boyle van ’t Hoff (BvH) relation, is that equilibrium cell water volume is proportional to the reciprocal of extracellular osmolality [8]. In response to changes in the solute concentration of the extracellular medium, water flows across the cell membrane toward osmotic equilibrium. The Boyle van ’t Hoff relation has been observed in many mammalian cell types, including chondrocytes [9] [10], endothelial cells [11], lymphocytes [12], and human islets [13] and HeLa cells [14]. However, the BvH relation does not account for cellular volume-regulatory mechanisms, including mechanosensitive channels, ion pumps, and other membrane transport pathways. Thus, although the BvH relation provides a useful description of equilibrium osmotic behaviour, it does not capture the time-dependent processes governing cellular volume regulation. Portions of this text were previously published as part of a thesis [15].

Under significant osmotic challenges, cell volume regulation plays a vital role in maintaining cellular homeostasis [16] [17]. Cells initiate volume regulatory mechanism when they swell or shrink in an anisosmotic medium. Accordingly, cells alter the activity of specific ion channels and transporters during anisosmotic stress to restore their volume toward the isosmotic state. For instance, when a cell becomes osmotically swollen, the process of regulatory volume decrease (RVD) causes it to release KCl, organic osmolytes, followed by water efflux, thereby restoring cell volume towards its initial value [18]. Conversely, osmotically shrunken cells gain NaCl and water to restore cell volume, a process known as regulatory volume increase (RVI) [19, 20]. Moreover, RVI has been associated with the activation of ion transport pathways, including the Na+/K+ pumps and Na+-K+-2Cl cotransporters (NKCC) in many cell types [18]. Therefore, it is necessary to develop a method for time-dependent measurements of cell volume changes to better understand the dynamics of cell volume regulation.

The physical characteristics of the nucleus differ from those of the cell, thus affecting its osmotically induced volumetric response. Osmotic responses are well understood at whole cell level; however, the osmotic properties of the nucleus are not well characterized. Transport between the nucleus and cytoplasm occurs through nuclear pore complexes, which regulate the exchange of molecules across the nuclear envelope [21]. Therefore, the nuclear envelope functions as a molecular sieve that allows the passage of ions and small solutes while restricting the diffusion of larger macromolecules [22, 23] [24]. Changes in extracellular osmolarity cause alterations in solute and water movement across the nuclear envelope. The relationship between nuclear volume and extracellular osmolarity has been shown to follow BvH-like behaviour under certain osmotic conditions, demonstrating that nuclear volume is osmotically responsive [1] [25] [26]. However simultaneous measurements of dynamic cell and nuclear volumetric responses remain limited. In articular chondrocytes, Finan et al found that the nucleus are linear osmometers under hyperosmotic conditions but exhibits a nonlinear relation under hyposmotic conditions [6]. These findings suggest that nuclear osmotic behaviour may differ from that of the whole cell, highlighting the importance of simultaneously measuring cell and nuclear volume changes.

Nucleoplasm differs from the cytoplasm in its organization as it contains macromolecular aggregates [27]. In turn, the concentration of macromolecules within the nucleus strongly influence the nucleus size by the excluded volume effect [28], where a molecule’s excluded volume corresponds to the area where its center is isolated from both the surrounding region and the space occupied by another molecule. Osmotically induced changes in cell volume influence macromolecule concentrations, aggregate formation, nuclear size, and chromatin condensation [27]. Changes in extracellular osmolality can alter intracellular macromolecular crowding, leading to changes in chromatin condensation and nucleocytoplasmic transport, and nuclear physical organization is known to have profound effects on cellular function [29]. For example, chromatin condensation is associated with apoptosis; accumulation of nuclear lamins A/C during stem cell differentiation is associated with increased nuclear stiffness; and alterations in nuclear size and morphology are characteristic of many cancer cells [30]. Nuclei have been examined in previous studies for their physical and biomechanical properties and are considered to be stiffer and more viscous than cytoplasm [2]. However, few studies have examined how osmotic stress affects cell and nuclear volume simultaneously, particularly in adherent cells, and how changes in macromolecular crowding contribute to the nuclear osmotic response.

Cryopreservation in a vitrified state has a considerable potential for long-term preservation of adherent cell monolayers due to avoidance of damage by ice formation [31] and could enable the ready supply of cells for pharmaco-toxicological studies. However, the high concentration of permeating solutes such as cryoprotective agents (CPAs) like DMSO or ethylene glycol required for vitrification is associated with osmotic damage from volume excursions during CPA addition and removal [32]. Osmotic pressure is exerted by the differences in the intracellular and extracellular solute concentrations across cell membranes and is considered as a major hurdle to successful cryopreservation of monolayers [17]. One approach to overcome these hurdles is to use modeling approaches to predict cell responses to osmotic stress [11, 33]. These models require the estimates of solute and water permeability parameters to design and optimize cryopreservation protocols. Measurement of these parameters in adherent cells remains challenging because of the limited availability of methods for directly measuring cell volume in adherent monolayers. The relative volume of adherent cell layers has been measured using several different techniques, including cell morphology, light scattering, and intracellular cation intensity changes [34], and calcein self-quenching in cultured endothelial cells [35] [36]. However, these indirect methods give an overall volume assessment, treating cells as a uniform block of space, and may not fully account for cell-specific morphology and subcellular organization. For example, polyploidy of hepatocytes can change the intracellular macromolecular crowding, potentially influencing interactions between the cytoplasm and nucleus and cellular transport [37] [38]. By precise quantitative estimation of cell and nuclear osmotic responses, we can understand the biophysical and biochemical mechanisms underlying cellular osmoregulation.

Here, we present the direct measurement of an individual cell and nuclear volume in monolayers in response to anisosmotic conditions. The overall goal of this study was to investigate osmotic behavior, volume regulation, and mechanical resistance to volume changes in cells and nuclei within a confluent monolayer. Studies have explored cellular osmotic responses to estimate biophysical, biochemical characteristics of hepatocytes isolated from rats, pigs, human [39] [40] [41] [42], however, these studies have primarily used indirect methods for the quantification of cell volume and associated biophysical parameters. For this study, we used human hepatoma HepG2 cells, as these cells are extensively used as an *in vitro* hepatic model, particularly in pharmacological and toxicological research [38, 43–46]. HepG2 cells show considerable nuclear heterogeneity, with a proportion of cells being binucleated or polyploid and frequently showing anisokaryotic nuclei [47–49]. Given the importance of the nucleus in determining cellular function and fate [50], understanding the effects of osmotic stress on both cells and nuclei simultaneously is important. We first hypothesized that HepG2 cells and nuclei function as ideal osmometers. Second, we hypothesized that the permeability of cells and nuclei to solutes and water can be accurately derived by precise measurement of volumetric excursions induced by osmotic stress. We investigated these hypotheses by measuring the dynamic and equilibrium volume of the cell and nuclei in response to changes in extracellular osmolarity using live 3D confocal microscopy for adherent HepG2 monolayers.

## 2. Materials and methods

### 2.1. Cell culture

The HepG2 human hepatoma cell line was obtained from the American Tissue Culture Collection (ATCC) using the clonal derivative (HepG2/C3A, Catalog no. CRL-10741). The cells were cultured in Eagle’s Minimum Essential Medium (MEM) supplemented with 10% FBS and 1.5 g/L sodium bicarbonate, non-essential amino acids, L-glutamine, and sodium pyruvate (Sigma-Aldrich, Canada) at 37 °C and 5% CO2. The media was changed every 48 hours during cell cultivation. When cells reached 60-70% confluency, they were dissociated with 0.25% Trypsin-EDTA solution and resuspended in fresh culture medium at an appropriate seeding density for continuous propagation. For experiments, HepG2 cells were seeded directly into 6-channel μ-Slide VI 0.4 (IBIDI, Wisconsin, USA) at a concentration of 1 x 106 cells/mL and resulted in a confluent cell monolayer.

### 2.2. Solutions

Hyper- and hyposmotic solutions containing only non-permeating solutes were prepared by adding NaCl stock solution (2 Osm/kg) or distilled water to the isosmotic (300 mOsm/kg) phosphate-buffered saline (PBS containing 0.905 mM Ca2+ and 0.492 mM Mg2+) to create solution osmolalities of 80, 145, 200, 600, 730, and 1215 mOsm/kg. For permeating solutes, a 1 mol/kg solution was made by adding dimethyl sulphoxide (DMSO), or ethylene glycol (EG) (ThermoFisher Scientific, Mississauga, ON, Canada) to isosmotic PBS. All osmolalities were measured using freezing point depression osmometer, Fiske 2400 multisample osmometer (Fiske Associates, Norwood, Massachusetts, USA).

### 2.3. Image acquisition

Cells were treated with calcein-AM (1.25 µg/mL) and Hoechst 33342 (1.0 µg/mL) solution in isosmotic (290 ± 5 mOsm/kg) PBS for 15 min in a 37 °C incubator. The dye-loaded cells were subsequently exposed to anisosmotic solutions of different osmolarity ranging from (80, 145, 200, 600, 730, 1215 mOsm/kg) and time-lapse images were acquired throughout the osmotic exposure. Cells were initially maintained in isosmotic medium which was carefully replaced with appropriate hypotonic and hypertonic solutions using a Pasteur pipette to minimize physical disruption of the monolayer during imaging.

Fluorescent imaging was performed using an inverted Zeiss LSM 880 confocal laser scanning microscope equipped with Airyscan using 40X /1.2W Korr FCSM27 objective. To acquire single labeled images of the cells, standard filter sets and laser lines were used in the imaging process. Calcein and Hoechst 33342 fluorescence were excited with the 490 nm and 350 nm laser lines, respectively, and the emission was assessed at wavelengths 520 nm and 460 nm, respectively. For volumetric imaging, *z*-stacks were acquired from the coverslip to beyond the apical surface of the cells to capture the entire cell volume within the region of interest (ROI). The thickness of the cells and the step size determine the number of 2D images per *z*-stack of the cells being imaged. Each optical section consisted of 512 x 512 pixels with a *z*-step size of 0.5-1 µm. Depending on cell thickness and the selected z-step size, approximately 20–25 optical sections were acquired per z-stack. Each optical section required approximately 0.18–0.21 s to acquire, resulting in a total acquisition time of approximately 4–6 s per complete 3D z-stack.

To get the baseline cell and nuclear volume under isosmotic conditions, 4–5 z-stacks were acquired before replacing the isosmotic medium with the respective anisosmotic solution. Time-lapse 3D imaging was then continued until the cells reached equilibrium with the applied solution. Next, to characterize cell and nuclear responses to permeating solutes, monolayers were similarly exposed to 1 mol/kg permeating solute solution of DMSO or EG. Cells were exposed to respective non-permeating or permeating anisosmotic solutions for 160 s under hyperosmotic conditions and 300 s under hyposmotic conditions. Following exposure, the anisosmotic solution was replaced with isosmotic medium, and imaging was continued for an additional 160 s to characterize volume recovery following restoration of isosmotic conditions. A total of 15-30 individual cells analyzed per treatment and each experiment was replicated twice for this study.

### 2.4. Image analysis

In this study, Hoechst 33342 labelled volume was directly used as the nuclear volume, while the calcein-labelled volume was used to define the total cell volume (**Fig 1A**). Nuclei were used as seed points for the identification and separation of individual cells, with the outer cell boundary segmented based on calcein fluorescence. Cytoplasmic volume was calculated by subtracting the Hoechst labelled nuclear volume from the corresponding calcein-defined total cell volume as follows:

Hoechst labelled volume= Nucleus Volume (*V*_nuc_),
Calcein labelled volume= Total Cell Volume (*V*_cell_),
Cytoplasmic Volume (*V*_cyto_) = Total cell volume (*V*_cell_) - Nucleus Volume (*V*_nuc_).

**Fig 1:**
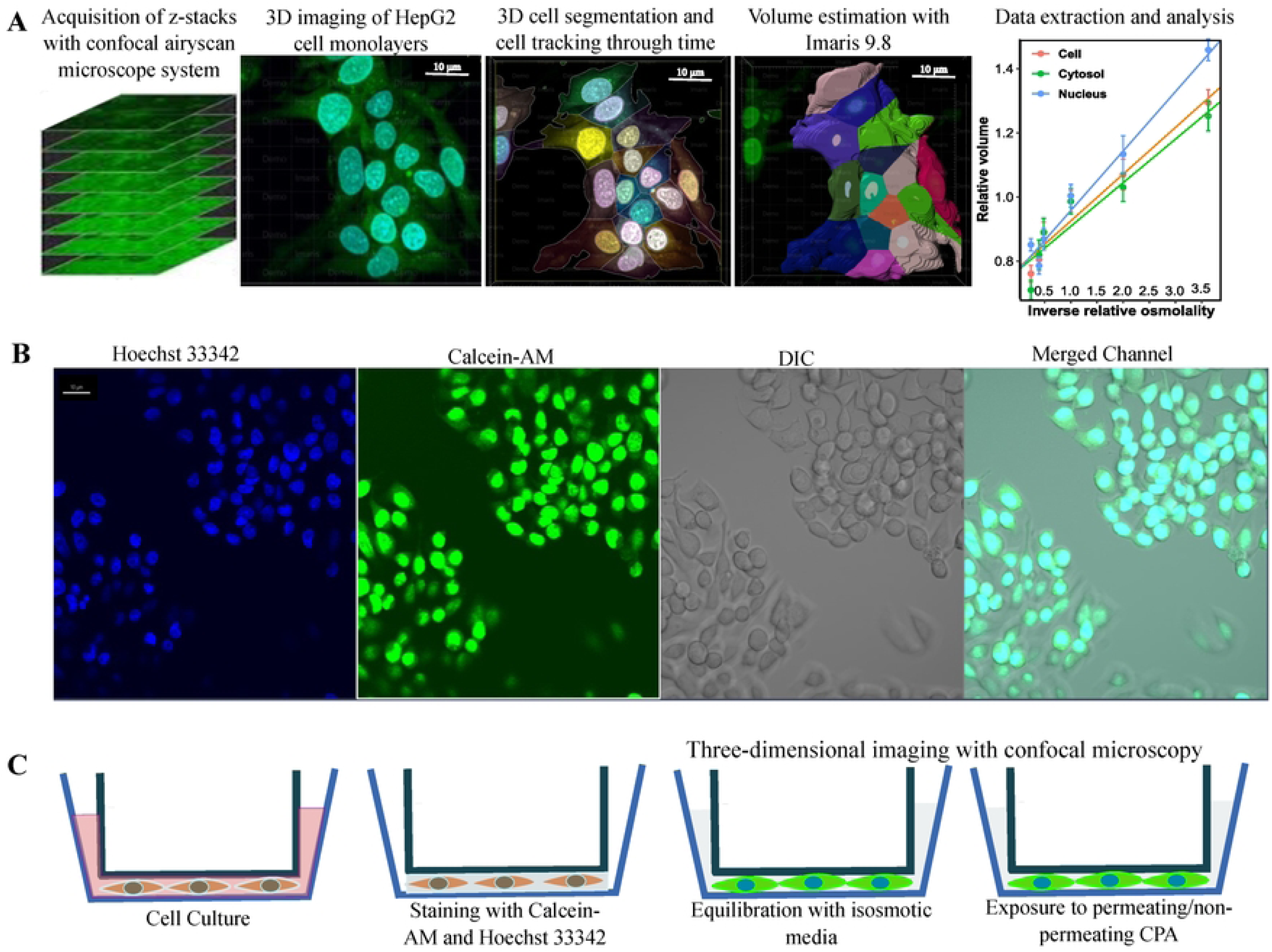
**A.** A sequential representation of image acquisition and analysis method for adherent HepG2 cells. **B.** Correlative fluorescent and brightfield micrographs of HepG2 cell monolayers. The first (left to right) channel (blue) shows nuclei, the second (green) shows the cell, the third (grayscale) shows the differential interference contrast (DIC) image for HepG2 monolayers, and the fourth represents the merged channels at 10X magnification. (**C**) An illustration of the experimental workflow for imaging and exposure of adherent HepG2 cells to permeating and non-permeating solutions.

Images were obtained after Airyscan processing using built-in software functions of ZEN 2.3 (blue edition) followed by segmentation and 3D reconstruction with image analysis software Imaris 9.0 (Bitplane Oxford-Instrument) for 3D visualization and data processing (**Fig 1B**). For segmentation, we adopted rigorous criteria considering pixel intensity, size, and shape to delineate cell and nuclear boundaries accurately. In the cell module of Imaris, segmentation was done by using the cell module, using nuclei as seed points and splitting cells based on seed points. Custom code in Mathematica was used to ensure nuclei were within the respective cell throughout time via nearest neighbor algorithm with a threshold radius equal to the mean cell radius, providing tracked cell-nucleus pairs throughout time. Additional filtering was done based on the intensity and quality of the fluorescence by applying smoothing and local contrast filtering. Bleb formation was observed as a normal physiological response in HepG2 cells during the hypo-osmotic challenge, and lysed cells were excluded from data analysis. Surfaces were reconstructed based on pixel grouping, and cells were tracked over time using high-performance motion algorithms. This method extracts time information, including cell, cytoplasm, and nuclear surface area, volume, diameter, sphericity, and spatial position. Finally, cell and nuclear segmentation was additionally assessed using the machine learning software Cellpose, with the Cellpose “cyto” model and nuclei as a channel to guide segmentation of the 3D images [51, 52].

### 2.5. Mathematical Modeling

Cell volume, *V*_cell_, consists of an osmotically active water volume, *W*_cell_, that responds to changes in extracellular osmolality, volume of solutes such as CPAs, and an osmotically inactive volume, 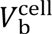, that includes cellular solids and bound water that does not readily participate in osmotic volume changes, such that 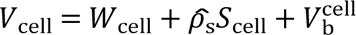, where *S*_cell_ is the moles of solutes in the cell and *ρ*_s_ is the molar volume. The cell is composed of the cytoplasm (*V*_cyto_) and nucleus (*V*_nuc_) such that *V*_cell_ = *V*_cyto_ + *V*_nuc_. Both cytoplasm and nucleus are themselves composed of osmotically active and inactive fractions in the forms 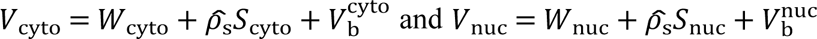 respectively. The Boyle van ’t Hoff (BvH) model is the most widely used model to determine the osmotically active water volume and the osmotically inactive (non-osmotic) volume of the cell [8]. The BvH model states that the intracellular water volume *W* can be determined from the Boyle van ’t Hoff relationship (Eq. 1) [53, 54] [55],

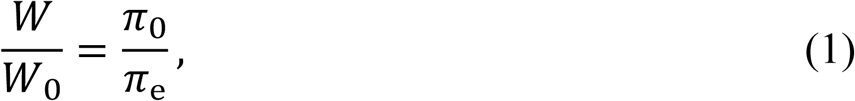

where, *π*_e_ and *π*_0_ are the extracellular and a reference osmolality (typically set to isotonic osmolality, *π*_iso_), respectively, and *W* and *W*_0_ are the equilibrium total cell water volume at osmolalities *π*_e_ and *π*_0_, respectively. Cells are called ideal osmometers if their volume at osmotic equilibrium varies linearly with the reciprocal of the osmolality of nonpermeable solutes in the external medium.

Here we wish to account for the transport of water and a permeating solute across both the cell and nuclear membranes into two compartments separated by semi-permeable membranes (**Our current** method directly fits for the cytoplasm and nuclear parameters and confined the search space to be within two orders of magnitude larger than literature values.

Fig 2). To do so, we extend the usual two parameter model by defining variables *W*_cyto_ and *W*_nuc_ for the volume of water in the cytoplasm and nucleus, respectively, *S*_cyto_ and *S*_nuc_ for the moles of permeating solute in the cytoplasm and nucleus, respectively. Thus, we defined a nested compartment 2P model that accounts for flux of water and solutes across the cell membrane but also across the nucleus membrane simultaneously. This mass-transfer equation (typically referred to as the two-parameter (2P) model) is based on Fick’s law that describes the flux of membrane-permeable solutes [56] [57]. In this case, we assume that passive transport of solutes is appropriate: on the scale of large osmotic shifts over short time periods, there is insufficient time to move a significant quantity of solutes via active transport that would not be captured in the phenomenological transport constants for water and solute. Thus, the water and solute fluxes can be written as

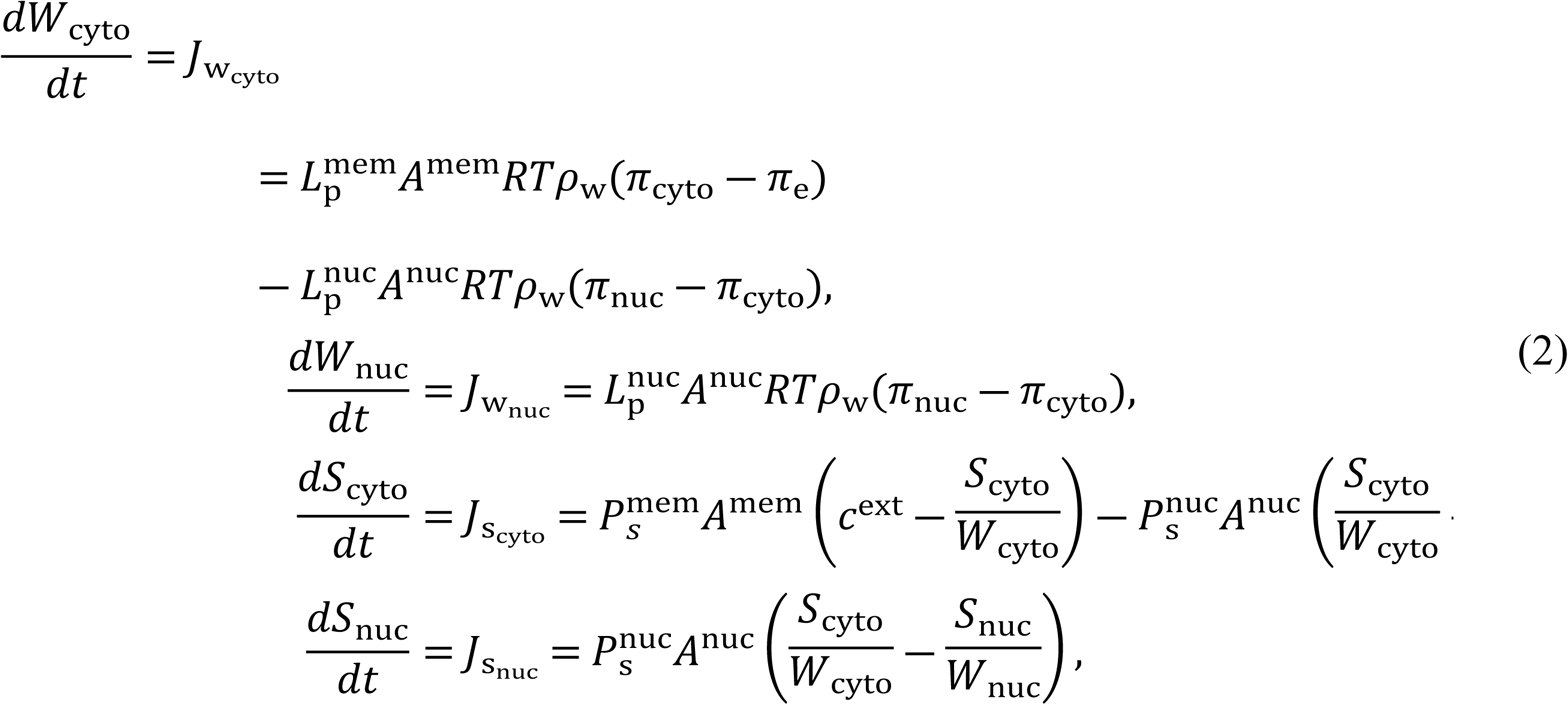

**Fig 2:**
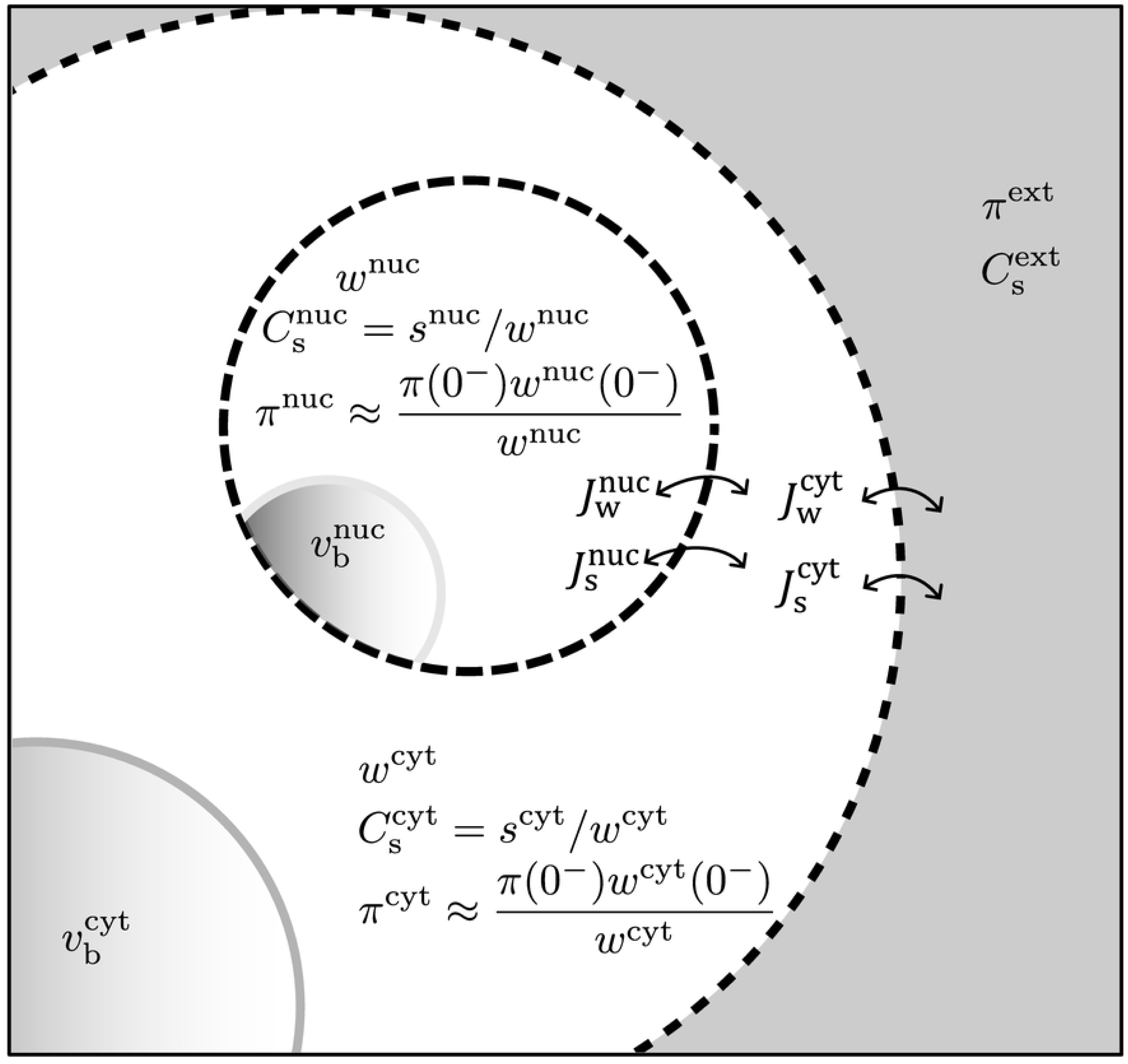
A schematic representation of double boundary two parameter model. The outer circle represents the cell membrane that separates the cytoplasm from extracellular fluid. The inner circle represents the nuclear envelope. Here, 0-indicates the initial values that are determined from the equilibrated value in isosmotic condition.

with initial conditions

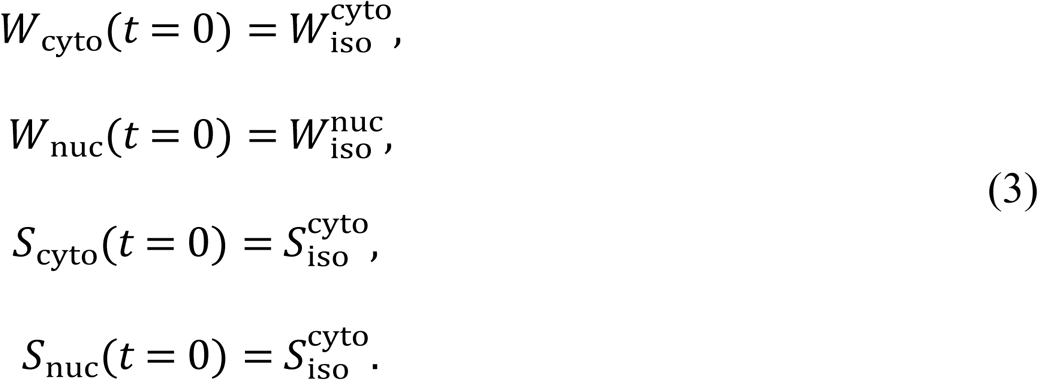

Here *π*_e_ is the extracelluar osmolality, *c*^ext^ is the extracellular concentration of permeating solutes, intracellular osmolalities are defined by 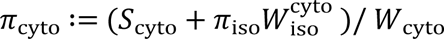, and 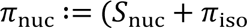 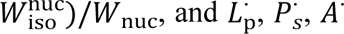 represent the hydraulic conductivity, permeating solute permeability, and surface area, respectively, of the cell membrane (*mem*) and nuclear membrane (*nuc)*. Notably, here we let *J* represent the flux into the compartment indicated by the subscript, not necessarily the flux across a specific membrane. Furthermore, we note the term 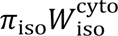 provides the intracellular (typically) non-permeating osmolytes *N* for the respective compartment. Finally, Eq. 2, as a representation of compartmental flux, can be used to recover the usual whole-cell transport model (cf. Benson [58], Abazari [59], etc) as

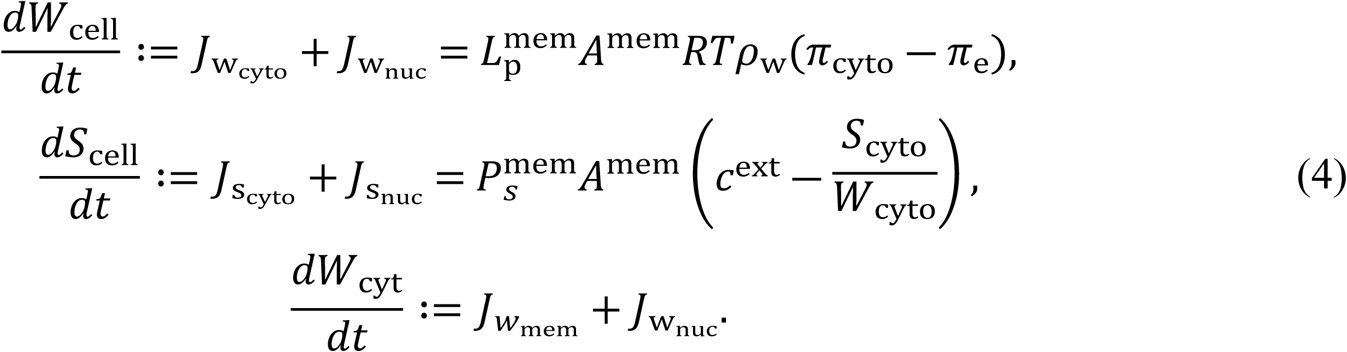

Equilibrium is reached when the net flux of water and solute across the cell membrane and nuclear membrane is zero and is satisfied when,

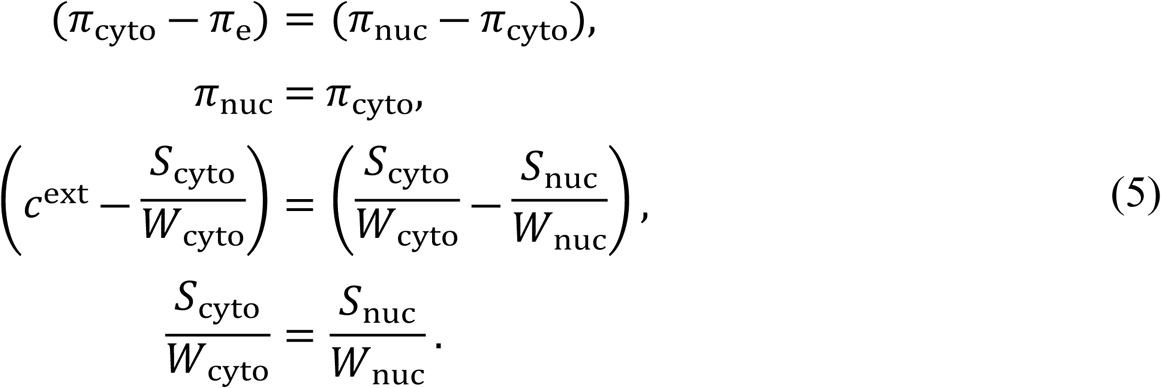

The equilibrated compartment volumes may be retrieved by recalling 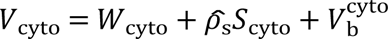 and 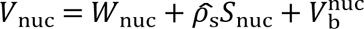 with the explicitly in the form written as,

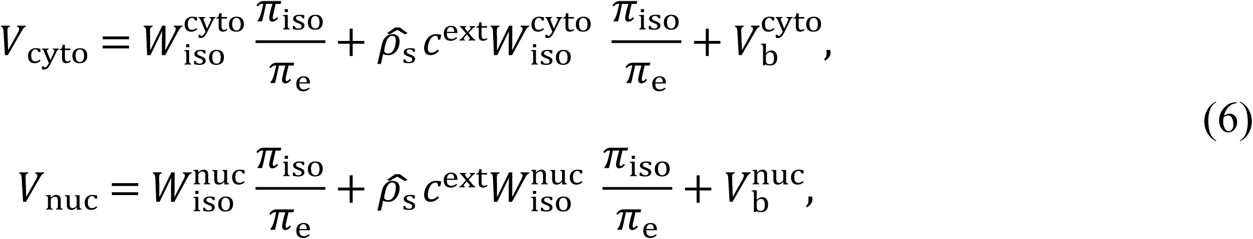

and the total equilibrated cell volume is similarly retrieved in the form

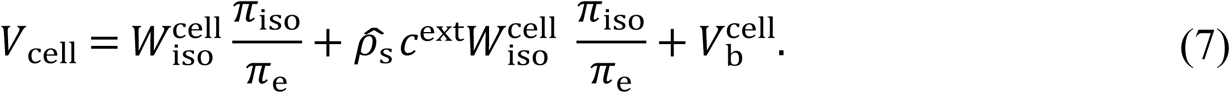

### 2.6. Statistical Analysis

We performed linear regression analysis for cell volume and inverse osmolarity for the Boyle van ‘t Hoff relation. All volume data were normalized with respect to their respective initial isosmotic volume. To test for autocorrelation in the output of a regression model, a Durbin-Watson statistic (DW) was used, and this score is reported with each figure. An ANCOVA was used to compare the slopes and intercepts (osmotically inactive volume) among the regression lines. Moreover, to find the differences among the treatments, a two-way ANOVA was conducted and t-test or a Tukey HSD was used to determine the differences among components i.e., cell, cytoplasm, or nucleus, respectively. All statistical analysis was conducted in R and Mathematica v14, with significance level set to P < 0.05.

For volume kinetics data, Eq. 2 represents a system of ordinary differential equations which was nondimensionalized and numerically solved in Mathematica v14 using *NDSolve*. To fit the 2P model to data we used Mathematica’s *NMinimize* function with the *Differential Evolution* method to find the minimum of the objective function (here the residual sum of squares, RSS) with respect to the cytoplasm and nucleus volumes throughout time. When fitting to volume data, Eq. 2 has 4 to 6 fitted parameters (represented as the vector *ϴ*) and two predicted volumes 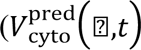 and 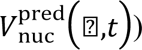 for a specific time, *t*. As the cytoplasmic volume and nuclear volume were significantly different and biased fitting for the cytoplasm volume, the volumes were normalized with respect to the isotonic volumes and objective function is defined as

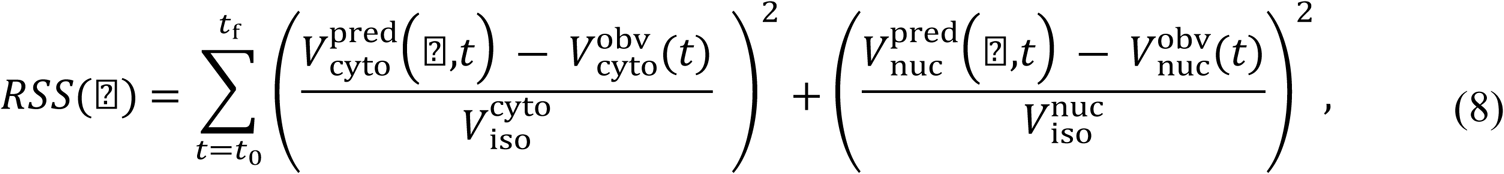

where the superscript “obv” refers to the respective observation at time *t*. Our current method directly fits for the cytoplasm and nuclear parameters and confined the search space to be within two orders of magnitude larger than literature values.

## 3. Results

### 3.1. Equilibrium Volumetric responses of the cell and nucleus

A sequence of 3D volume images of adherent HepG2 cells after exposure to hypo and hyper-osmotic media is shown in **Fig 3**. The images are presented as 2D images and stained with calcein-AM and Hoechst 33324 for the cell, cytoplasm, and nuclei. **Fig. 3A** displays the time-series images of HepG2 cells during hyposmotic exposure, where it can be observed that cells start to swell and form blebs. The blebs are marked in the image. **Fig. 3B** shows the time-series images of HepG2 cells during hyperosmotic exposure, where the cytoplasm and nuclei shrink resulting in a change in cell and nuclei morphology.

**Fig 3:**
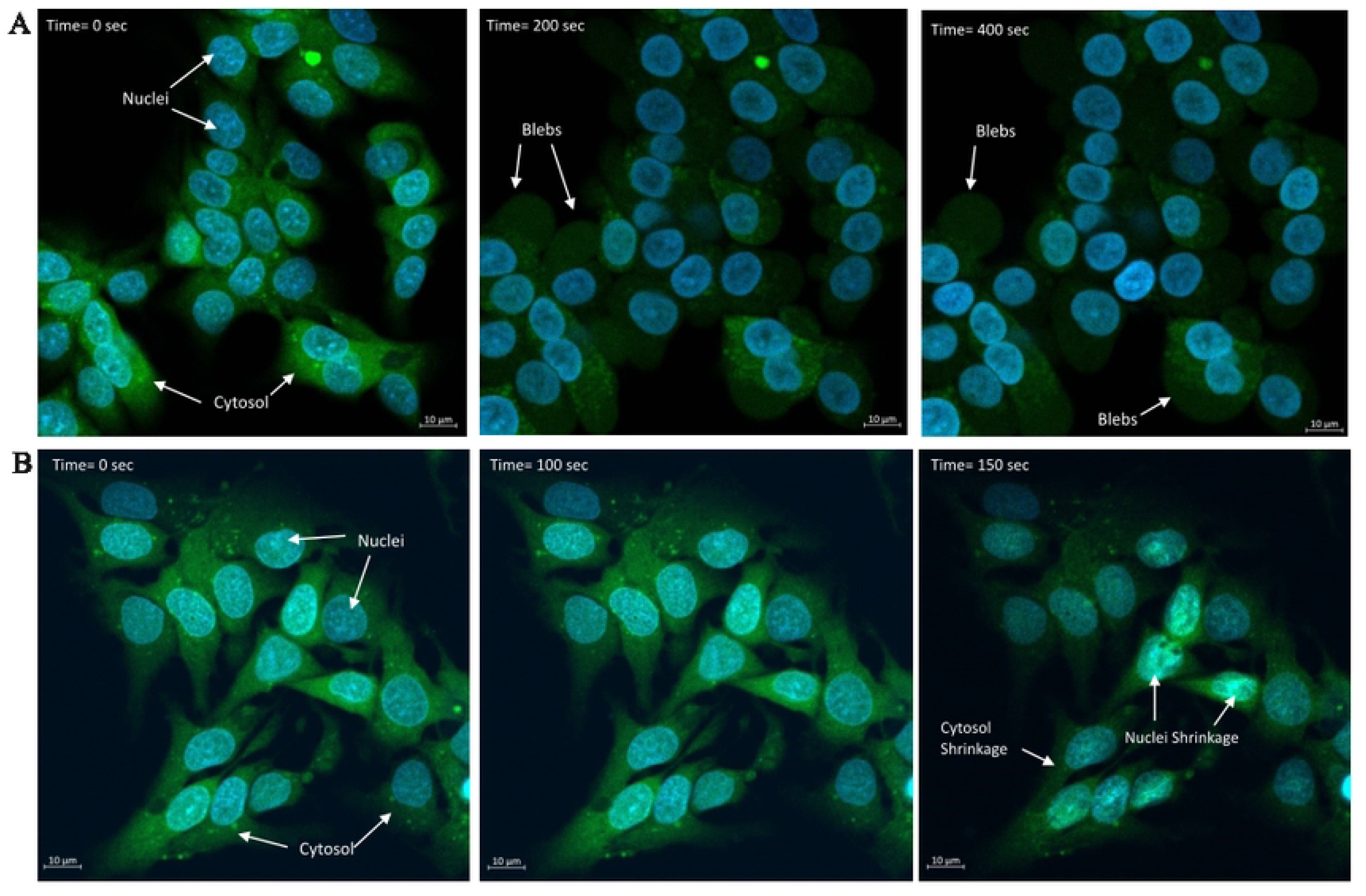
A sequence of 3D volume images of adherent HepG2 cells after exposure to hypo and hyper-osmotic media are presented as 2D images, stained with calcein-AM and Hoechst 33324 for cell, cytoplasm, and nuclei. (**A**) represents the times series images of HepG2 cells during hyposmotic exposure (80 mOsm/kg). It is shown that cells start swelling and blebbing, and bleb can be seen as labelled in images (**Supplementary video Fig 3A_S1**). (**B**) represents the times series images of HepG2 cells during hyperosmotic exposure (1215 mOsm/kg). Hypertonic stress results in shrinking of the cytoplasm and nuclei, and this can be observed as a change in cell and nuclei morphology (**Supplementary video Fig 3B_S2**).

We investigated volumetric equilibration behaviour HepG2 monolayers exposed to various osmolarities at room temperature (21 °C). A linear relationship was fit to inverse normalized osmolarity and cell volume over a range of anisosmotic conditions tested (80 to 1215 mOsm/kg) (**Fig. 4A**). The osmotically inactive volume was found by extrapolating normalized volume data in the BvH plot to infinite osmolality. Osmotically inactive volumes represent the minimum volume of tightly packed cellular constituents. To understand the relationship between relative volume and relative osmolality, we conducted linear regressions (BvH) for cytoplasm, nuclei, and total cell volumes over the full osmotic range. We found the same solid volume fraction (0.77) for cytoplasm, nuclei, and the cell given by the intercept (1/π = 0) of the linear regression (**Fig. 4A**). An analysis of covariance (ANCOVA) to compare the regression lines, suggests that slopes of the regression lines for each component are significantly different (F_2, 3530_ = 380, P < 0.001). Interestingly, in the hypotonic region, a higher positive slope (0.18) was found for nuclei, which was significantly different (P < 0.001) from cytoplasm (0.13) and the cell (0.140). This indicates that the cytoplasm preferentially loses water (likely through loss of ions) in comparison to the nucleus. The Durbin-Watson *d* statistic indicates a positive autocorrelation among the residuals for total cytoplasm, cell and nuclear volumes with values of *d*=1.126, 1.076 and 2.552, respectively, and a critical region bounded between 1.869 (*d*_L_) and 1.899 (*d_U_*) at 5% significance. Finally, a two-way ANOVA showed a significant effect of cell structure (F=8.816, P < 0.001), with significant effect of osmotic (hypo and hyperosmotic) stress (F= 2.610, P < 0.001), and a significant interaction between the structure (cell, cytoplasm, and nucleus) and osmotic stress (F= 0.212, P > 0.05).

**Fig 4:**
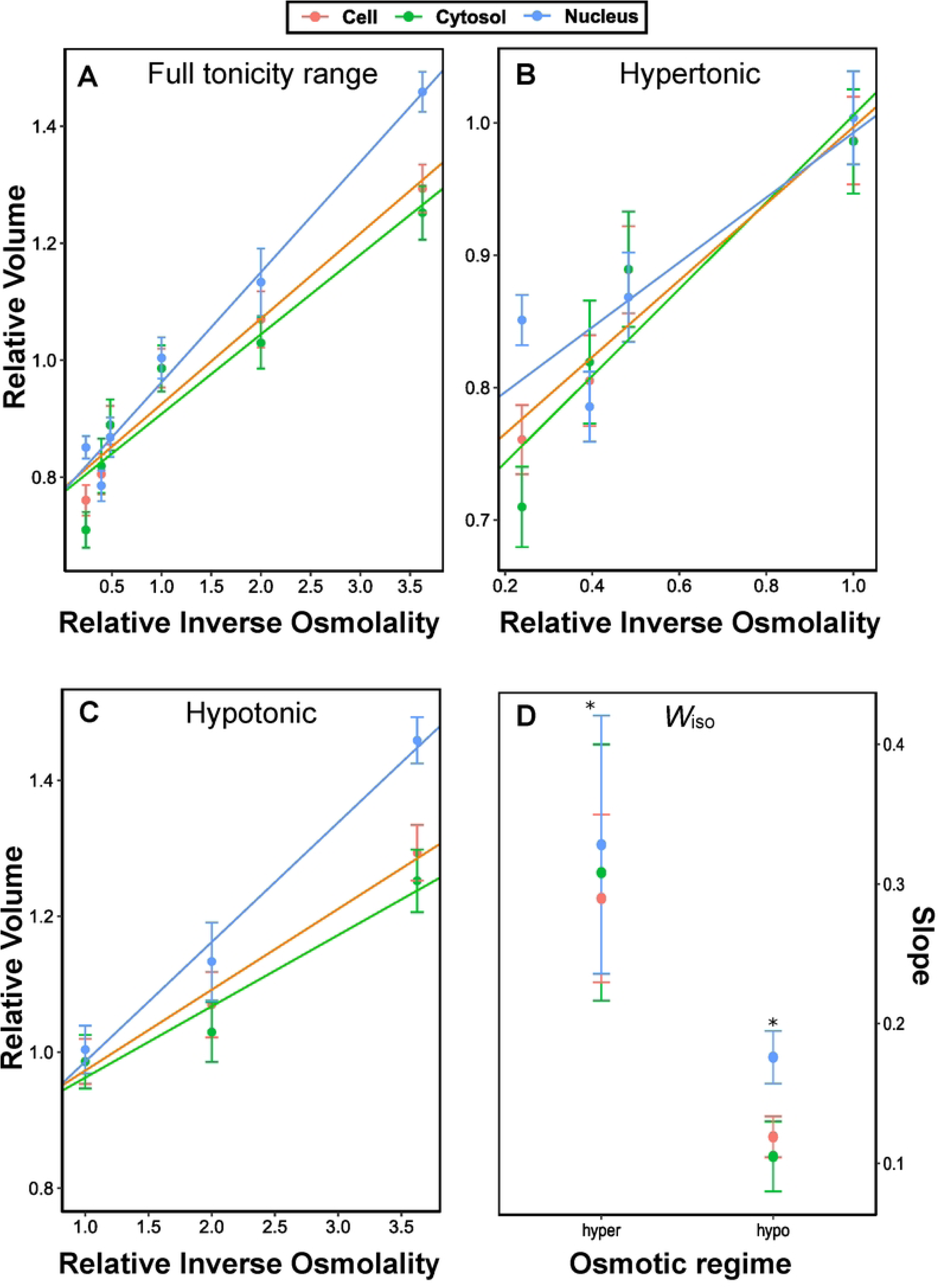
Boyle van ‘t Hoff plots for equilibrium cell, cytoplasm, and nuclear volumes for the entire osmotic range **(A)** and for the separate hyper **(B)** and hypotonic **(C)** ranges for the HepG2 monolayers. Volumes are normalized to their respective initial isosmotic volume, and dots and error bars represent mean ± standard error of the mean. Slopes for cell, cytoplasm, and nuclear volume are represented with orange, green and blue colors, respectively. In **(A)**, R2 values for total cytoplasm, cell and nuclear volumes were 0.8979, 0.9513, and 0.9776 respectively with DW scores to be at 1.126, 1.076 and 2.552 respectively. In (**B**), for the hypertonic region only, R2 values for total cytoplasm, cell and nuclear volumes were 0.8647, 0.9209, and 0.8785 respectively with DW scores of 1.892, 2.571 and 2.567, respectively. In (**C**), for the hypotonic region only, R2 values for total cytoplasm, cell and nuclear volumes were 0.9463, 0.9851, and 0.9886 respectively with DW scores of 2.962 for all components. **(D)** Comparison of regression analysis slopes (representing *W*_iso_ for the cell and each compartment) under different osmotic conditions, showing nuclei have higher slopes under all conditions. Symbol (*) shows statistically significant differences among each osmotic condition (P < 0.05), and dots and error bars represent mean ± standard error of the mean. Between 15 and 30 individual cells were analyzed per treatment.

For comparison, BvH plots were separated into two plots for hypo and hypertonic ranges. In each case, a linear regression of the data was performed to allow comparisons of the equilibrium volume (**Fig 4B**). In the hypotonic range, the slope of the nuclei was significantly steeper (0.17) than that of the cytoplasm (0.10) and whole cell (0.12) (ANCOVA, F_2,1200_ = 124, P < 0.001), suggesting more relative nuclear expansion in hypotonic regions. Further, we found no positive autocorrelation among the residuals for their respective linear regression models, with Durbin-Watson *d* statistics to be at 2.962 for all components, and a critical region bounded between 1.791 (*d*_L_) and 1.831 (*d_U_*) at 5% significance.

In the hypertonic region, slope of the nuclei (0.24) is significantly less (ANCOVA, F_2, 1730_ = 112, P < 0.001) than the cytoplasm (0.33), and the cell (0.29) (**Fig 4C**), with an inactive nuclear volume fraction of 0.78. Interestingly, cell, cytoplasm and nuclei showed different inactive volume fraction in hypertonic conditions. Moreover, we found no autocorrelation among the output of their respective linear regression models, with Durbin-Watson *d* statistics for total cytoplasm, cell, and nuclear volumes to be at 1.892, 2.571 and 2.567 respectively, with a critical region bounded by 1.802 (*d*_L_) and 1.848 (*d_U_*) at 5% significance. Altogether, comparison of all regression analysis slopes among different osmotic conditions by Tukey-HSD test showed that nuclei have significantly higher slopes under all anisotonic conditions (P < 0.05) as shown in **Fig 4D**. Therefore, a correlation analysis was conducted to test the relationship between cell and nuclear volume in non-permeating solutes. In hypotonic conditions, there was no significant correlation between nuclear and cell volumes, but in hypertonic conditions this correlation was significant (P < 0.05) (Fig 5).

**Fig 5:**
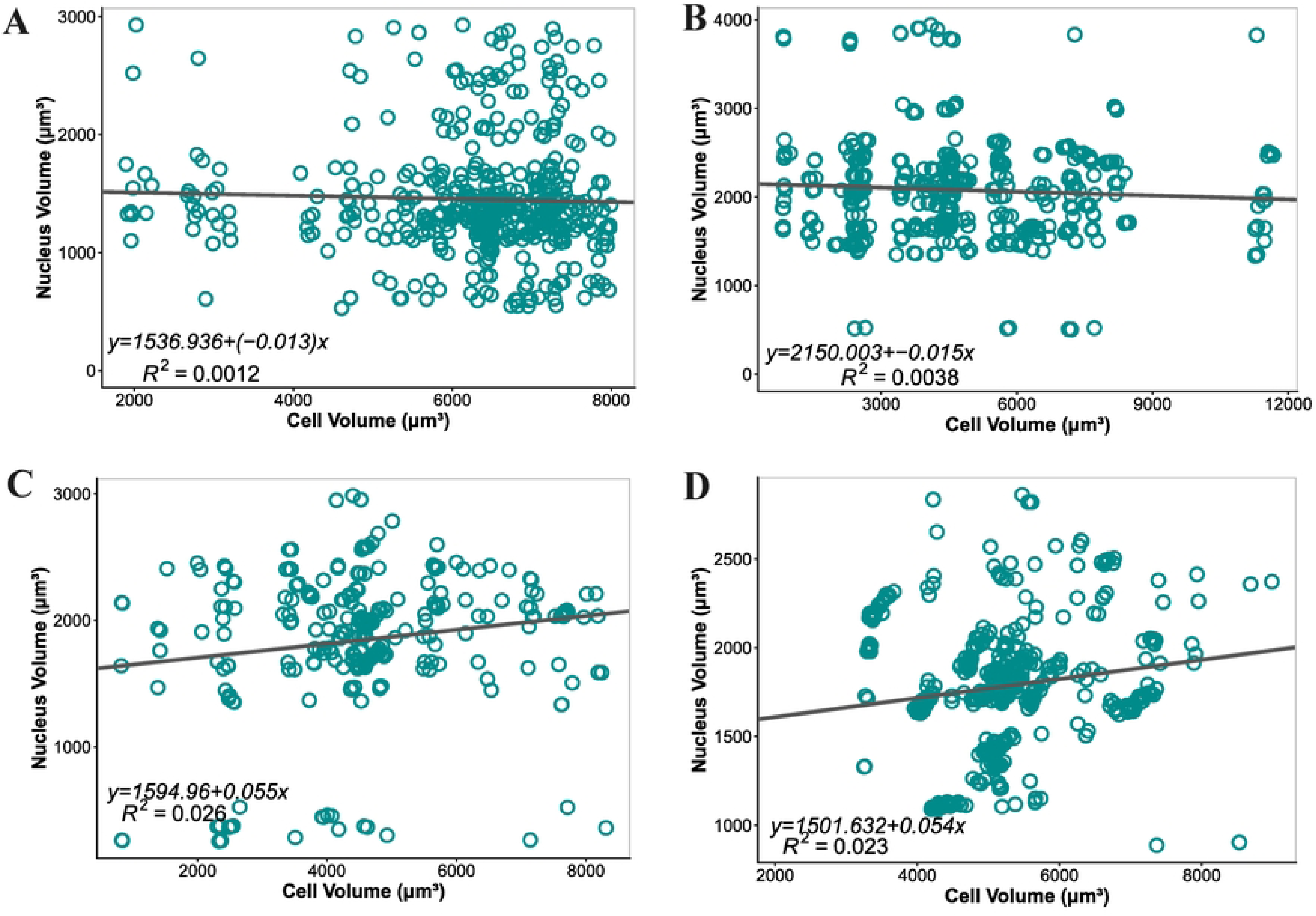
Correlation between cell and nuclear volumes in different anisotonic conditions. Panels (**A-D**) represent anisotonic solutions of 80, 200, 730, and 1215 mOsm/kg respectively. In the hypotonic region, nuclei undergo more expansion compared to the whole cell, therefore, no correlation was found between nuclear and cell volumes for 80 mOsm/kg (n = 676) and for 200 mOsm/kg (n = 300), as shown in panels **A-B**, respectively. A significant (P < 0.05) positive correlation was found under hyperosmotic conditions for 730 mOsm/kg (n = 336) and for 1215 mOsm/kg (n = 400) as illustrated in panels **C-D**, respectively. Each dot represents an individual cell measurement while in equilibration in anisotonic media and black lines are best-fit linear regressions with best fit equations and coefficient of determination are shown.

### 3.2. Estimation of water and solute permeability parameters

To investigate if the cells osmoregulate during anisotonic challenge, cells were exposed and returned to initial isotonic conditions with cell-nuclei pairs tracked throughout time (**Fig 6**). We individually tracked 125 cell-nuclei pairs. Restricting to datasets with all positive volumes yielded 110 tracked pairs, more than 90% of expected observations (properly segmented). The average isotonic cell volume (n = 110) was 5334 ± 239 um3 with an average surface area of 2705 ± 86 μm2. The average isotonic relative nuclear volume 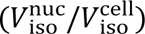 was 0.258 ± 0.011. Upon return to initial isotonic conditions there is a general trend of loss in cell volume for both hyper- and hypotonic treatments (**Fig. 6C-E**). Cells appeared to equilibrate to hypertonic exposures by an initial rapid decrease in volume within the first 30 s of exposure, followed by a slow decrease in volume thereafter with some cells exhibiting a change to a mild increase in volume (**Fig. 6C-E**). For hypertonic exposures, a multiple linear regression model describing volume after initial fitting with respect to cellID, osmolality, and time (restricted from 60 to 200 s), showed a significant (P < 0.05) decrease in relative cell, cytoplasm, and nuclear volume (**Table 2**). To check return volume, we normalized the cell and the compartments with respect to each respective isotonic point and tested significant difference using the Signed Ranked Test , with respect to 1, their normalized isotonic reference point. The Bonferroni adjusted alpha value is α_adj_ = 0.05/18 = 0.00278 (3 tests for each of the 6 treatments). We found that after hypertonic exposure, normalized cell and cytoplasmic volumes do not significantly differ from their initial isotonic volume (P > α_adj_, **Fig. 6E**), while noting the p-values are consistently low (P < 0.10) reflecting the observed volume trends in **Fig. 6C-E**. Exposure to 600 and 736 mOsm resulted in normalized nucleus volumes being significantly lower (P < α_adj_) than isotonic volumes after exposure. After exposure to hypotonic solution of 80 mOsm, the nucleus remained significantly enlarged (P < α_adj_) while the cytoplasm and overall cell normalized volumes significantly decreased (P < α_adj_) (**Fig. 6C-E**).

**Fig 6:**
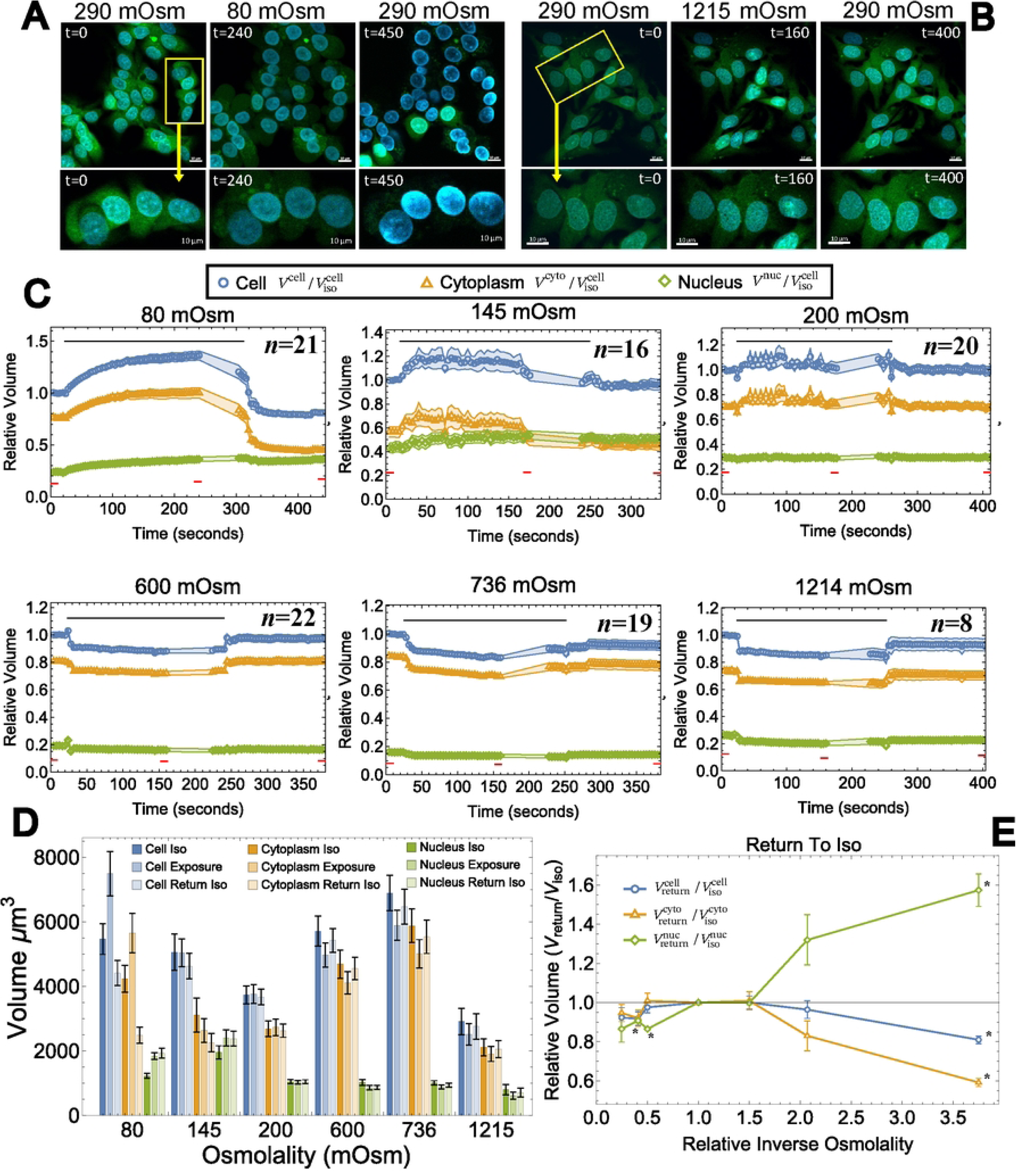
Monolayers of HepG2 cells are exposed to anisotonic conditions and return to isotonic conditions. (**A-B)** A representative figure of HepG2 cells exposed to a hypotonic solution of 80 mOsm and 1215 mOsm with the cytoplasm and nucleus shown before, during, and after osmotic challenge. **(C)** Relative cell, cytoplasm, and nucleus volumes are plotted throughout time. Symbols represent mean and bands represent SEM. Tracked cell-nucleus pairs are normalized with respect to their respective isotonic cell volume (first average volume of first 16 s). Red lines indicate the time frame used to obtain mean volumes used in panels D and E. Osmotic challenge is applied as indicated by the black bars. Cells are returned to isotonic conditions (300 mOsm). **(D)** A bar chart of average cell, cytoplasm, and nucleus volumes. Error bars represent SEM. Iso, exposure, and return iso volumes are the mean volumes for the timeframes indicated in red in Panel C. **(E).** Return to isotonic plot shows the normalized cell, cytoplasm, and nucleus volumes after return to isotonic conditions (return to iso volume normalized with respect to isotonic volume for the respective tracked cell-nucleus pair). Open symbols represent mean values, and error bars represent SEM. The * symbol represents a significant difference from normalized isotonic volume using a Spearman Ranked test and an adjusted alpha value of 0.05/18 = 0.00278, where 18 is the number of tests performed

**Table 1:** A description of the parameters and values used for two parameter-double boundary model fitting of adherent HepG2 cells volumetric responses for various anisosmotic and CPA solutions.

| Parameter | Description | Units | Value |
| --- | --- | --- | --- |
| $W$ | Intracellular water volume | $\mu\text{m}^3$ | variable |
| $W_{\text{iso}}^x$ | Water volume in x at isosmotic condition | $\mu\text{m}^3$ | variable |
| $\pi_x$ | Osmolality of x | $\text{mol kg}^{-1}$ | variable |
| $\pi_{\text{iso}}$ | Isosmotic osmolality | $\text{mol kg}^{-1}$ | 0.300 |
| $V_x$ | Volume of x | $\mu\text{m}^3$ | variable |
| $W_x$ | Osmotically active volume in x | $\mu\text{m}^3$ | variable |
| $V_b^x$ | Osmotically inactive volume in x | $\mu\text{m}^3$ | variable |
| $s$ | Normalized moles of permeating solute | N/A | variable |
| $S_x$ | Moles of permeating solute in x | mol | variable |
| $t$ | Incubation duration | min | variable |
| $T$ | Temperature | K | 295.15 |
| $R$ | Universal gas constant | $\mu\text{m}^{-3} \text{ atm}^{-1} \text{ K}^{-1} \text{ mol}^{-1}$ | $8.206 \times 10^{13}$ |
| $A$ | Cell surface area | $\mu\text{m}^2$ | specified |
| $v$ | Non-dimensional cell volume | $\mu\text{m}^3$ | variable |
| $\rho_w$ | Density of water/PBS | $\text{kg L}^{-1}$ | 1.00 |
| $\hat{\rho}_s$ | Molar volume of CPA | $\text{mL mol}^{-1}$ | 71.0 (DMSO)<br>55.7 (EG) |
| $c^{\text{ext}}$ | The extracellular concentration of permeating solutes | mol | variable |
| $P_s^x$ | Solute permeability of the x membrane | $\mu\text{m}/\text{min}$ | parameter |
| $L_p^x$ | Hydraulic permeability of the x membrane | $\mu\text{m}/\text{min}/\text{atm}$ | parameter |
| $J_w^x$ | Water volume flux into x | $\mu\text{m}^3 \text{ s}^{-1}$ | variable |
| $J_s^x$ | Solute volume flux into x | $\mu\text{m}^3 \text{ s}^{-1}$ | variable |
| cyto | Cytoplasm (cell minus nucleus) |  | Param. label |
| nuc | Nucleus |  | Param. label |
| cell | Whole cell |  | Param. label |
| mem | Cell membrane |  | Param. label |

**Table 2.** Parameter table for the loss of relative volume throughout time (s) for each compartment. The parameters are for the multiple linear regression with respect to osmolality (nominal), CellID (nominal) and time (60 s to 120 s). There are a total of 1221 observations, 49 unique cell-IDs and 3 hypertonic osmolalities. The p-values represent the probability that the respective parameter is equal to zero (no change in relative volume). While cells were normalized with respect to their own isotonic volumes, we note that the mean isotonic cell volume is 5,334 ± 239 μm3.

|  | *Relative volume*<br>*Loss (sec<sup>-1</sup>)* | *Standard Error* | *t-Statistic* | *P-Value* |
| --- | --- | --- | --- | --- |
| $V_{cell}/V_{iso}^{cell}$ | $-3.16 \times 10^{-4}$ | $7.47 \times 10^{-6}$ | -42.2911 | $<10^{-5}$ |
| $V_{cyto}/V_{iso}^{cell}$ | $-2.58 \times 10^{-4}$ | $7.64 \times 10^{-6}$ | -33.7687 | $<10^{-5}$ |
| $V_{nuc}/V_{iso}^{cell}$ | $-0.581 \times 10^{-4}$ | $1.96 \times 10^{-6}$ | -29.7146 | $<10^{-5}$ |

We estimated the water and solute permeability parameters for both the cell and nuclei by using water and solute flux equations (Eq. 2). We used the segmented surface area associated with each tracked cell during fitting and asserted that approximately half of the surface area was in contact with the media. We fit Eq. 2 with respect to each individual cell-nucleus pair with each fit presented in supplementary materials (**Fig. S3**). Upon visual inspection, some datasets (∼18) were phenomenologically unrealistic with unlikely jumps in volume possibly due to error in segmentation and/or tracking (**Fig. S3**). A model fit with unrealistic data provides equally unrealistic behaviour, typically with extreme parameter values. Unrealistic data may be pruned via thresholding methods based on quantifiable characteristics of the dataset with respect to the distribution across all datasets. To help mitigate pruning bias, we present the median of the fitted parameters here in **Table 3**. For completeness, we also performed a pruning method and found the mean values are similarly to simply using all datasets and then investigating the median behaviour indicating an agreement between the two methods; pruning/analysis is found in accompanying code. Using the median parameters provides a good predictive model for simultaneously estimating the relative volumes of the cell, cytoplasm, and nucleus across 6 anisotonic treatments with a total of 4 fitted parameters (**Fig**. **7A**). We present the correlated distribution of solid fractions for the cytoplasm and the nucleus alongside the hydraulic conductivity in **Fig. 7B**. Pruning the solid fractions data by selecting values that are in the range of 0.01 to 0.99 removes 28 datasets with the remaining datasets having a significant positive correlation of 0.321 between the cytoplasmic and nuclear solids fraction using a Spearman Rank test (n = 82, *r*_s_ = 0.302, P = 0.006) that is evident in **Fig. 7B**. There is no clear correlation between the cell and nuclear membrane permeability distribution.

**Fig 7:**
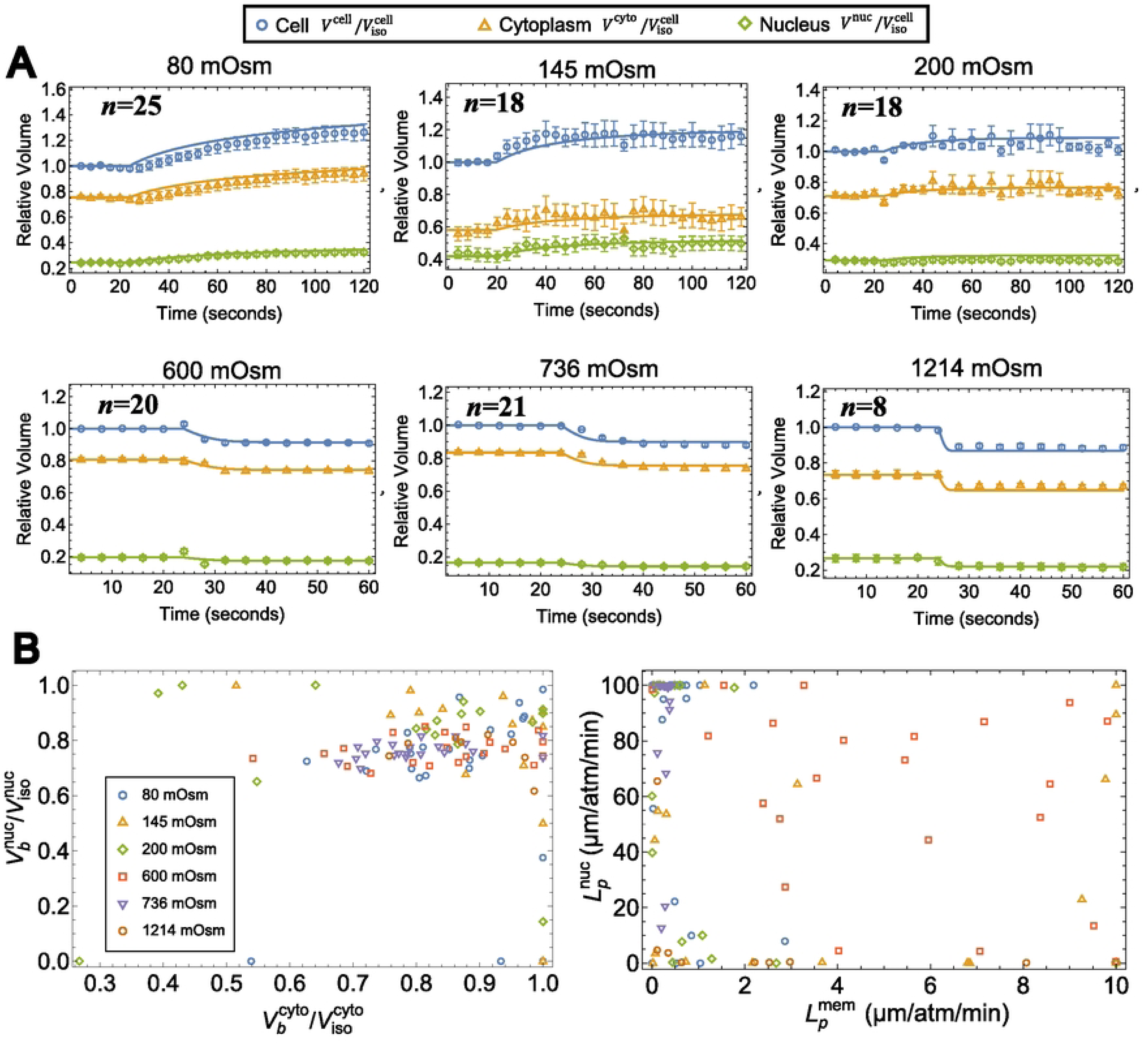
Presentation of the efficacy of Eq. 2 to model anisotonic exposures (xPBS). **(A).** The relative cell, cytoplasm, and nucleus volume responses for tracked cell-nucleus pairs exposed to various anisotonic solutions. All volumes are normalized with respect to their own respective isotonic cell volume (mean volume of the first 16 s in isotonic solution). Symbols represent the mean and the error bars represent SEM. The solid lines represent the predicted volumes by Eq 2 using the median of the fitted parameters at the individual level across all osmotic conditions (n=110; see **Table 3**). **(B)** The distributions of relative osmotically inactive fractions for the nucleus and cytoplasm are presented, alongside the hydraulic conductivity for the cell and nuclear membranes

**Table 3:**
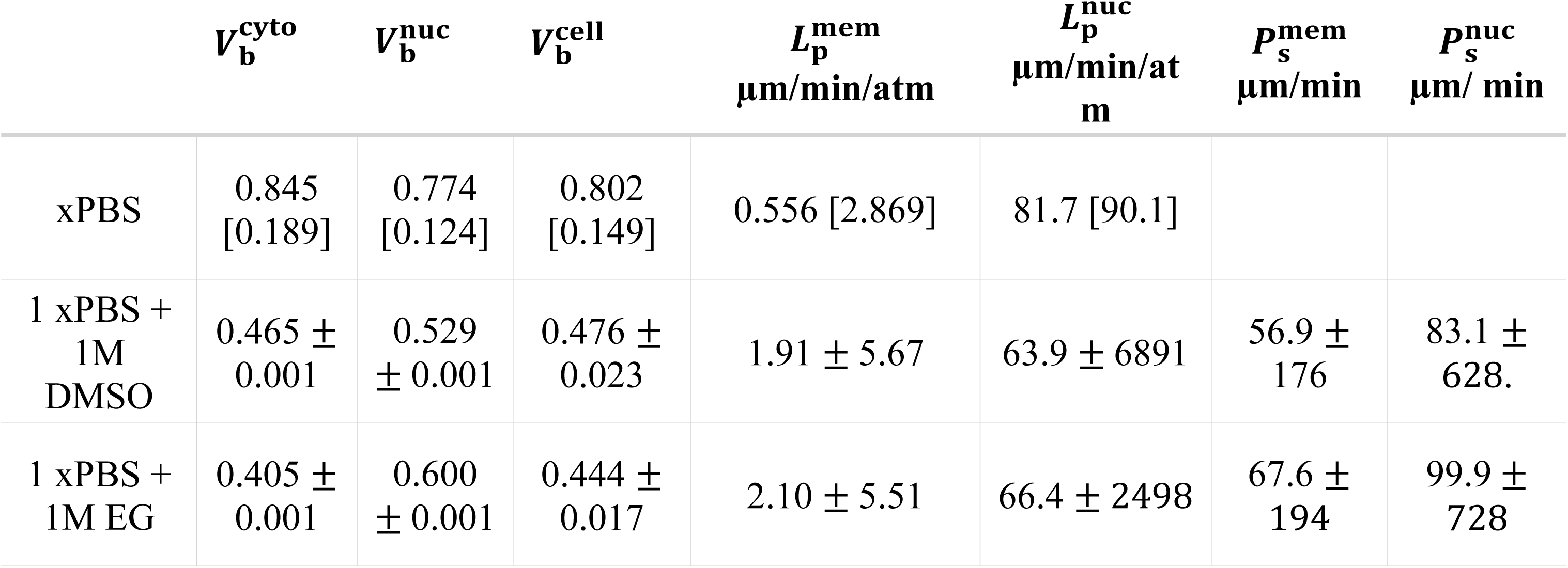
Membrane characteristics for the cell and nucleus under different anisosmotic conditions with and without dimethyl sulfoxide (DMSO) and ethylene glycol (EG). Experiments were conducted at room temperature. Anisotonic data is for tracked cell-nucleus pairs (n = 110). For DMSO and EG exposures, Eq. 2 was modified to include a heavy step function that changes intracellular osmolytes solved via Eqs. 6-7 given the observed equilibration volumes of the respective compartment. Eq. 2 with DMSO exposure was fit to a total of 1496 observations across four replicates and an average of 36 cells per time point. Eq. 2 with EG exposure was fit to 1505 observations across two replicates with an average of 35 cells per time point. IQR is the interquartile range and error ±SE. The parameter 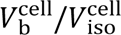 is not fit but calculated from the cytoplasm and nucleus osmotically inactive values and respective compartmental volumes; it is presented here for completeness.

The nuclear membrane hydraulic conductivity was of particular interest in this study. Using the same pruned fits, the average nuclear membrane hydraulic conductivity was 159 ± 22 times higher than the cell membrane hydraulic conductivity for the same cell-nuclear pair. This ratio grows to 1.6 ⨉ 104x using all pair fits. Notably, 43 out of 110 nuclear hydraulic conductivity fits were >99 µm/min/atm and near to the upper bound set as 100 µm/min/atm. This high hydraulic conductivity means the nucleus equilibrates with respect to the cytoplasm much faster than the change in cytoplasmic concentration change, and the model is relatively insensitive to changes in 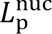 when 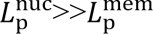. Figure S4 shows how nuclear volume becomes insensitive to changes in 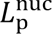 when 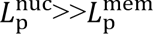. Higher boundaries were investigated but did not significantly improve the fits while reducing the probability of finding the confined global minimum (data not shown). Instead, it is possible the high fitted 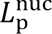 parameters emerge from not modelling ion flux across the nuclear membrane, which would reduce time to equilibration and reduce the effective hydraulic conductivity. Such a model may be represented by Eq. 2 with 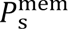 = 0 and allowing *S* to represent electrically neutral nuclear permeable free solutes and net neutral ion flux such as K+ and Cl-pairs). This model requires fitting permeability of the nuclear membrane intracellular free solutes, and the ratio of free to fixed intracellular solutes. Indeed, a similar model has been recently presented by Hoffmann and colleagues [60]. We fit Eq. 2 with intracellular osmolyte flux to each individually tracked cell-nucleus pair and compared the AIC between the two models. Contrary to our expectations, this osmolyte flux model did not result in a reduction of 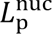 and only saw an improved (lower) AIC for 2 out of the 110 of the fitted datasets with a worse (higher) AIC for the remaining 108 datasets in comparison to the initial model. The model presented in [60] also included nuclear membrane tension resulting in a hydrostatic gradient across the nuclear membrane (see also [6]). We included a nuclear hydrostatic pressure gradient with osmolyte flux into Eq. 2 but again did not find any notable improvements in terms of AIC with 7 out of 110 found to have a lower AIC in comparison to Eq. 2 (analysis found in the supplementary code).

We investigated the effects of the permeable solutes DMSO and EG on volumetric kinetics of HepG2 monolayers by exposing them to 1xPBS supplemented with 1 mol/L DMSO or EG. The cells and compartments appeared to shrink lower than expected from the fitted model with anisotonic xPBS data (**Fig. 8**). This is possibly due to the implicit assumption of the cells behaving as ideal osmometers in the construction of Eq. 2. Furthermore, the cells appeared to equilibrate within 20 to 40 s of exposure, with a lower equilibration volume than expected with Eqs. 6-7 (**Fig. 8**). We used a simple multiple linear regression model for observations between 30 and 70 s and found the slope with respect to time was not significantly different from zero−indicating equilibration had occurred (**Table S1**). While there was a consistently lower observed equilibrated volume than expected (**Table S2**), this difference was not found to be significantly different (**Table S1**). The possible loss of volume during shrinking is in line with the significant loss of volume for hypertonic exposures (**Fig. 6**; **Table 2**).

**Fig 8:**
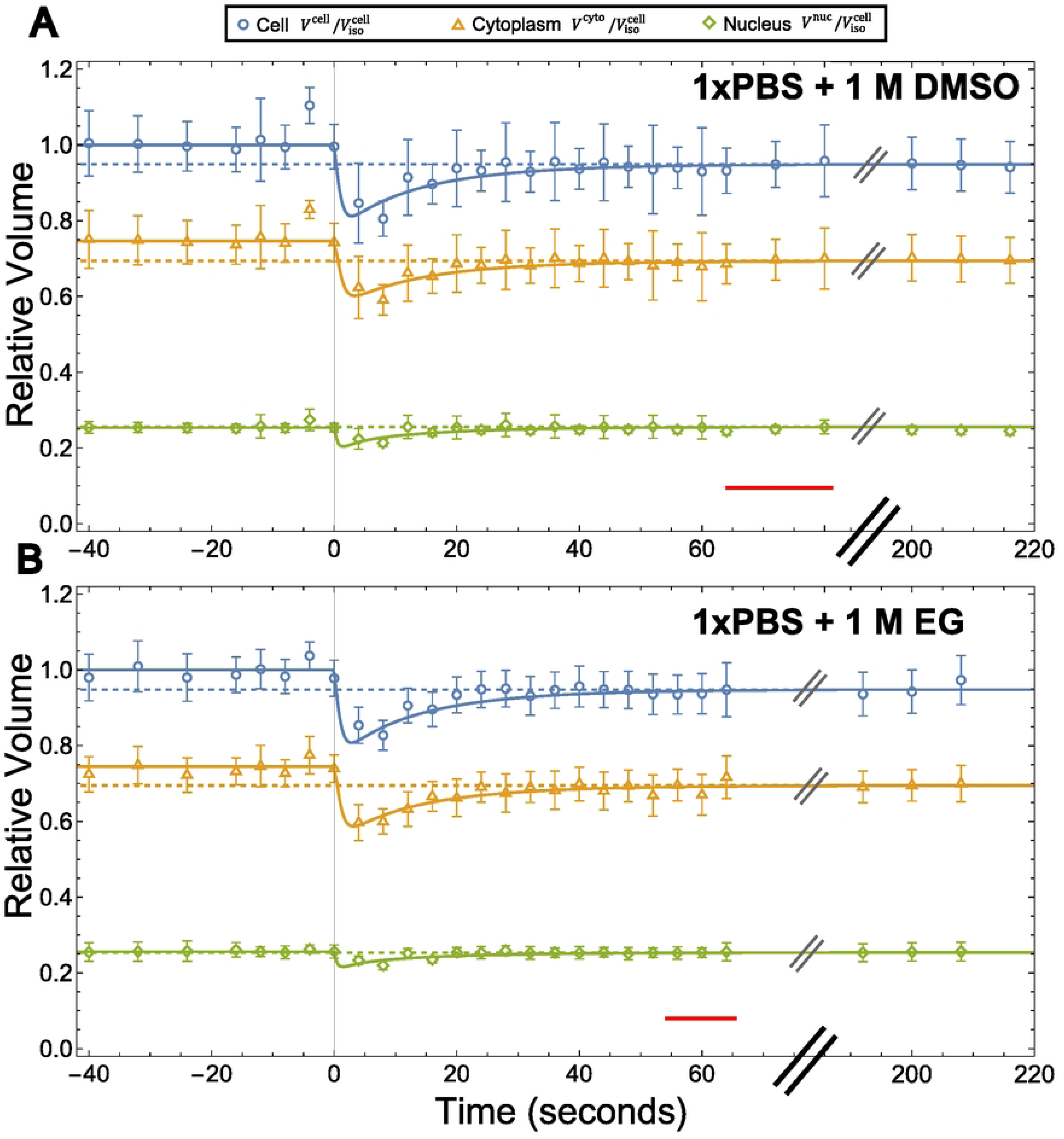
Volumetric response of HepG2 monolayer cell, cytoplasm and nuclear compartments exposed to 1xPBS with 1 mol/L dimethyl sulfoxide (DMSO) or ethylene glycol (EG) at room temperature. Compartmental volume is normalized with respect to initial isotonic cell volume at the population level. Solid lines represent Eq 2 with fit compartmental solids fraction and with intracellular osmolytes modelled via a heavy step function with the shift solved via Eqs. 6-7 given the observed equilibration volumes of the respective compartment indicated by the red bar. Dashed lines represent equilibrated volume of the respective cell/compartment. Symbols represent means and error bars represent SEM. **(A)** Volumetric response to DMSO exposure with a total of 1496 observations across four replicates and an average of 36 cells per time point. **(B)** Volumetric response to EG exposure with 1505 observations across two replicates with an average of 35 cells per time point.

Using the insights provided by the simple multiple linear regression model, we investigated modifications of Eq. 2. First, we allowed the osmotically inactive fraction of each compartment to be equal to the anisotonic xPBS exposures, or to be fit freely. Second, we used a heavy step function to model a shift in intracellular osmolytes of each compartment by solving Eqs. 6-7 with respect to observed equilibration volumes. We first fit the models without a temporal minimum volume requirement and obtain parameters that predicted minimum volume to be reached within a second of CPA exposure with very high fitted membrane permeabilities (only ever limited to set upper boundary of 102 μm/atm/min or 102 μm/min. That is, the model fit for the minimum RSS but at the expense of a reasonable minimum volume, which phenomenologically appears to occur ∼4 to 12 s (**Fig. 8**).

We therefore performed the fits again with the time constraint for the minimal volume to fall within 3 to 16 s. This resulted in more realistic parameters for the associated minimum RSS. In both cases, we found that the best performing model (in terms of AIC) used freely fit solid fractions and used the heavy step function to account for a loss in osmolytes/volume (**Table S3;** unbounded model comparisons are provided in the supplementary code). The time constrained model fits are provided in **Table 3** with the associated model plotted in **Fig. 8**.

## 4. Discussion

### 4.1. BvH relationship: HepG2 Monolayers are not ideal osmometers

Here, we presented a quantitative method for the simultaneous estimation of cell and nuclear volumetric responses under significant osmotic challenge. Human hepatoblastoma HepG2 cells were chosen to study the osmotic behavior of cells in the form of monolayers. We first examined the cell and nuclear osmotic behavior over a range of osmolalitites from 80 to 1215 mOsm/kg and found that equilibrium cell and nuclear volumetric responses of HepG2 cell monolayers are not ideal osmometers and not described by Boyle van ’t Hoff relation as indicated by the presence of autocorrelation and significant change between osmotic regimes (**Fig. 4**). The nonideal-osmometer behaviour of HepG2 cell monolayers sets HepG2 cells apart from other adherent cells that behave as ideal osmometers [61] [62] [63] [64]. Instead, HepG2 cells retain their nonideal osmometer behaviour while cultured as monolayers and during suspension [65].

We found that using the BvH relation provided an osmotically inactive fraction of 0.77 for each compartment normalized with respect to its own isotonic volume (**Fig 4**). When fitting Eq. 2 to anisotonic exposures of xPBS the osmotically inactive fraction was 0.85 for the cytoplasm and 0.77 for the nucleus and 0.80 for the cell (**Table 2**). When fitting in the presence of DMSO and EG, the osmotically inactive fractions decrease to 0.47 - 0.53 and 0.40 - 0.60 for the cytoplasm and nucleus respectively, and 0.44 - 0.48 for the cell. Li and colleagues [66] exposed lightly adhered (but still rounded in morphology) HepG2 cells to varying concentrations of PBS and to 1xPBS + 1.5 mol/L DMSO in a single step or in three steps. They recorded a 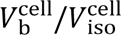 of 0.507 ± 0.064 in anisotonic solution, and 0.485 ± 0.087 for single step addition of 1.5 mol/L DMSO. During multistep loading of DMSO cell solids estimate dropped to 0.294 ± 0.094. Notably, in our past work [65], we found that suspended HepG2 cells do not behave as ideal osmometers, as assumed by Li et al. [66]. Indeed, the ideal osmometer assumption provided a 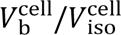 of 0.52 ± 0.02 and fit the data poorly, but using a non-ideal model (such as a leak model or turgor model), or just hypertonic data, resulted in a lower 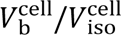 of 0.18 ± 0.04 [65]. In the current study, the hypertonic BvH data (**Fig. 2B**) do not provide a 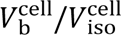 closer to the estimates using CPA exposure (**Table 3**) than fitting to the entire dataset (**Fig. 2A**).

It is worth considering the range of cell solids estimates provided by the current analysis. We found the mean isotonic volume of monolayers was 5334 ± 239 μm3. The corresponding dimensional osmotically inactive volume of the cell ranges from 2347 ± 105 μm3 to 4267 ± 191 μm3. Our past study on suspended HepG2 cells had isotonic volume of 1769 ± 59 μm3 and an osmotically inactive volume fraction of 0.18 ± 0.04 [65], with the corresponding dimensional volume being 318 ± 72 μm3. The transition from monolayer to suspended form is associated with a three-fold loss of total volume and surprisingly only retaining 7% to 14% of cell solids. The significant loss of cellular components may cause cells to be more susceptible to osmotic damage as suspended HepG2 cells were found to be much more osmotically sensitive than plated HepG2 cells [67].

### 4.2. Regulatory volume decrease and increase

A possible explanation for why the hypertonic data does not provide a good estimate of osmotically inactive fraction and the apparent loss of 86% to 93% of cell solids during the transition from monolayer to suspended is osmoregulation capability in hypertonic conditions when the cell is plated. We observed an initial decrease in cell volume during hypertonic exposure, followed by a less intense but significant decrease in volume thereafter (**Fig. 6**; **Table 2**). The rate of cell volume loss was 101 μm3/min (**Table 2**) and was dominated by loss of volume from the cytoplasm (**Fig 6C**). This rate of change in water volume may provide an estimate of osmolyte *N* (primarily K+ and Cl-) efflux by assuming that after the initial change in volume then *π*_e_≅*π*_i_ = *N*/(*ρ*_w_*W*), which means the flux of osmolytes is given by *dN*/*dt* = *π*_e_*ρ*_w_ *dW*/*dt*. Given *π*_e_ = 0.736 mol/kg then a water loss of 101 μm3/min corresponds to a solute flux value of 7.43×10-14 mol/min for typically nonpermeating solute such as K+ and Cl-. This efflux larger than the expected efflux via ion pumps of ∼1.81×10-15 mol/min [61] to ∼1×10−17 mol/min [65, 68], but is remarkably on the same order of magnitude we found for passive osmolyte leakage during regulatory volume decrease for suspended HepG2 cell in hypotonic conditions with ∼1×10−14 mol/min using the turgor-leak model and ∼4×10−13 mol/min using the *π*_e_≅*π*_i_ approximation [65]. For clarity, “osmolyte leakage” stems from the pump and leak concept, whereby there is a net passive flux of ions across the membrane (such as Cl-) [69] [70, 71]. Modes of leakage primarily include gated and non-gated and cotransport ion channels [68, 69, 71], and possibly during non-selective transmembrane gap formation due to chemical (such as DMSO exposure) [72] [73] [74] or temperature changes [75] [76, 77], all on top of the small but persistent lipid membrane diffusion [70]. If osmolyte leakage occurs passively, then solutes must flow down their concentration gradient. In hypotonic conditions the concentration gradient allows osmolytes to flow out of the cell, allowing for simplified models that group osmolytes into a single electrically neutral species [60, 65] [78] however in hypertonic conditions the reverse is expected.

The loss of ions during hypertonic exposure may be explained by a relatively higher conductance of K+ than Na+ across the membrane through opening of ion channels. That is, both K+ and Na+ flow down their concentration gradients (out of the cell and into the cell respectively), but the initial net efflux of K+ is larger than Na+ net influx. Indeed, Wehner and colleagues [61] found that HepG2 monolayer conducted both K+ and Na+ at a ratio of 2:1 and showed that whole cell conductance was 2.9 ± 0.3 nS in isotonic conditions but increased to 17.4 ± 1.2 nS during hypertonic exposure (isotonic salt media + 100 mOsm sucrose). Furthermore, they found that K+/Na+ ATPase was not a driving force behind regulatory volume increase/decrease. Instead, an increase in passive ion conductance, even at the expense of an increase of K+ conductance via opening of nonselective ion channels, occurred during osmotic challenge. We found a similar loss of volume (likely due to net efflux of osmolytes) in mouse embryonic stem cells after exposure to DMSO, chilling (−4 °C), or latrunculin A [*Under Review*]. Each of these treatments were hypothesized to cause a flux of calcium into the cell and/or result in the release of intracellular calcium stores, which in turn activated ion channels and allows for a net efflux of K+, and Cl-in isotonic conditions and nonselective ion flux in hypertonic conditions. Christmann et al. present data showing that chilling of HepG2 cells in isotonic conditions results in a decrease of cellular water content, indicating HepG2 cells may have similar regulatory volume mechanisms as mouse embryonic stem cells [79]. Indeed, a net loss of K+ and Cl-aligns with the observed loss of cell volume during hypertonic challenge and the loss of cell volume upon return to isotonic conditions (**Fig 6E**) and during equilibration to CPAs (**Fig. 7**). A longer equilibration in time is likely required to see significant regulatory volume increase to occur with the slower influx of Na+.The observation of loss of cell volume in hypertonic conditions is distinct from what is observed and predicted by Casula and colleagues [78]. They show that a loss of volume after exposure may be due to mechanosensitive ion channels opening during return to isotonic conditions, not during hypertonic exposure such as is the case in our current study. Importantly, our observation of hypertonic volume loss requires a mathematical representation of unique osmolyte/ion species within and outside the cell. To rationally predict regulatory volume decrease/increase and the return to isotonic conditions of HepG2 cells in anisosmotic conditions one must include electro-diffusion, mechanical tension, ion channels, and nuclear pores into a nested mechano-electro-osmotic model. We leave the development of such a model for future work.

### 4.3. Nuclear volume response to anisotonic conditions

Several studies report that nuclear membranes allow small (30-60 kDa) molecules to pass freely by passive diffusion from the cytoplasm to the nucleus and vice versa [21, 80, 81]. These molecules are considered small in the context of nuclear pore complex permeability. However, larger molecules (larger than 60kDa) have also been reported to permeate the nuclear pore complex (NPCs) *in vivo* over a period of many minutes to hours[82, 83]. Similarly, a previous study has revealed that small molecules diffuse across the nuclear membrane at such a fast rate that they are considered freely permeable to nuclear envelopes[84].

Previously, it was suggested that eukaryotes exhibit a strong correlation between cell volume and nucleus volume [85, 86], i.e., changes in the extracellular osmolarity will affect the nucleocytoplasmic transport that in turn will change the size of the cell and nucleus[87, 88]. When we investigated the relationship between cell and nuclear volume by conducting a correlation analysis, we found that nuclei undergo greater relative expansion compared to the cell during hypotonic challenge; therefore, a negative correlation was found between nuclear and cell volume (**Fig** 5). However, a positive correlation was observed between nuclear and cell volume in the presence of hypertonic conditions. In comparison, previous studies have examined the volume of the cell and nucleus during the growth cycle and in transformed cell lines[85, 86], but have not investigated it with respect to osmotic stress during hypotonic swelling and hypertonic shrinking. Strikingly upon returning the cell to isotonic conditions, the nuclear volume is larger than the initial isotonic nuclear volume (**Fig. 6E**). The increase in nuclear volume may result from chromatid decondensing, DNA fragmentation, and histone decomposition, increasing osmolyte concentration and stiffening the nuclear lamina. This is in accordance with the observations from Finan and colleagues, who found an increase in stiffness of nuclear lamina during hyposmotic exposure [6, 89]. A schematic explanation for these behaviours is shown in **Fig 9**. Interestingly, when exposed for 10 minutes to pure water, the nuclei of Hela, RPE-1, N2a, U-2 OS, COS-7, and HEK 293 all decrease below their initial isotonic volume [90]. It is possible that holding HepG2 cells in pure water for 10 minutes may result in a similar reduction of nuclear volume as the loss of intracellular osmolytes would likely be extreme, possibly offsetting mechanisms increasing the nucleus size. Finally, the results of our study are consistent with those reported by Kim and colleagues for fibroblasts [26], as they reported volume regulation and shape bifurcation in nucleus. However, further study is required to investigate the effect of osmotic stress on the physiological functions of the cell.

**Fig 9:**
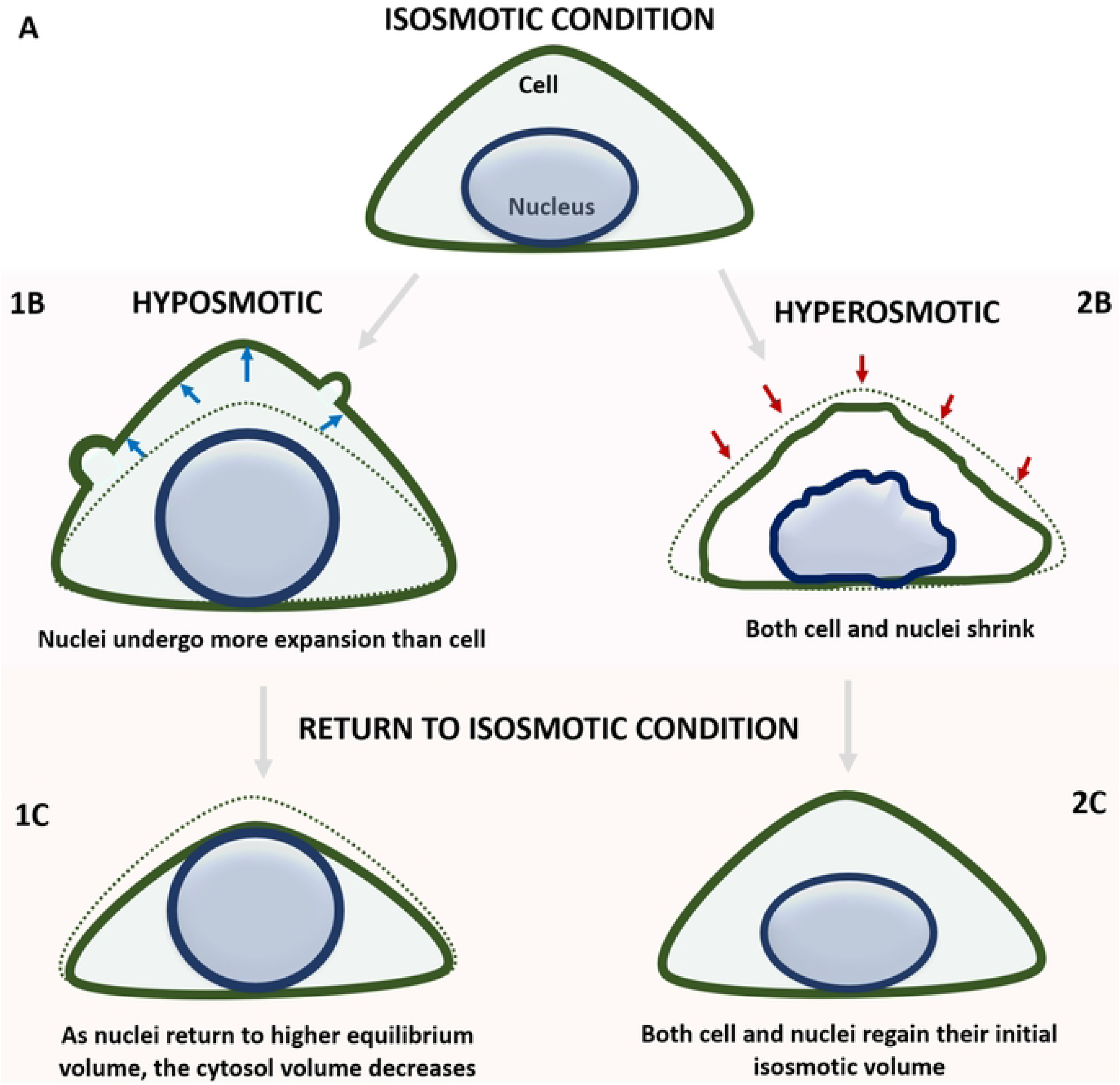
Experimental evaluation of the cell and nuclear volumetric responses in HepG2 monolayers (A). The outer cell membrane separates the cytosol from the extracellular environment and prevents large solutes from diffusing freely. The inner circle represents the nuclear envelope, that isolates the cytoplasm from the nucleoplasm and allows the diffusion of small molecules, but not large ones. Both cell and nuclei osmoregulate when exposed to anisosmotic media. In hyposmotic media (1B), both cell and nuclei start swelling, however, nuclei undergo more expansion than the cell. Whereas, both cell and nuclei undergo volume loss during hyperosmotic stress (2B). During hyposmotic stress, outward osmotic pressure is shown with blue arrows, whereas inward osmotic pressure is represented with red arrows during hyperosmotic stress. Alternatively, when cells are returned to isosmotic media after osmotic stress, nuclei return to higher inactive volume fraction after hyposmotic stress, with the cytosol lost its osmolytes/ions, returning to less volume than initial isosmotic volume (1C). Whereas, after hyperosmotic stress, both cell and nuclei return to their initial isosmotic volume (2C).

### 4.4. Membrane Permeabilities

Our analysis of anisotonic exposure and return to isosmotic conditions shows that HepG2 cells are not ideal osmometers. Although limited, Eq. 2 is still useful to describe short (30 to 90 s) exposure durations in anisotonic conditions (**Fig. 7**) but will likely become increasingly less applicable in longer time periods or multistep protocols (**Fig. 6**). In the presence of permeable solutes, modifying Eq. 2 to account for a loss of intracellular osmolytes improved equilibration volume prediction (**Fig. 8**). Interestingly, Li and colleagues [66] found single step protocols did not perform well to predict volumetric response to permeating cryoprotectant loading. Predicting more accurate multiple loading and unloading steps will likely require a more sophisticated model of ion flux model. However, is interesting to note that equilibration happens within 20 to 40 s HepG2 monolayers (**Fig. 8**), much faster than observed for suspended HepG2 cells that are lightly adhered (but still spherical) that do not reach volumetric equilibrium within 500 s of exposure to 1xPBS + 1.5 M Me2SO [66]. Reflecting this difference in equilibration time are how the monolayers have fitted *P*_s_ values a magnitude higher (57 ± 176 μm/min DMSO, 66 ± 194 μm/min EG; **Table 3**) than suspended HepG2 cells (3.6 ± 1.1 μm/min for single step and 7.1 ± 2.4 μm/min for multistep) [66]. Furthermore, given the rounded morphology in [66], we extracted cell diameters from their Fig. 1c and obtained a mean volume of 1705 ± 189 μm3 and surface area of 682 ± 50 μm2, assuming a spherical geometry. For their spherical cells, the surface area to volume ratio is 0.40 ± 0.05 μm-1 while for the monolayer cells in the current study the ratio is 0.51 ± 0.03 μm-1. This means that the monolayers have a higher surface area to volume ratio accounting for size, making permeation of cryoprotectants relatively faster. For hydraulic conductivity of the nucleus, past work has either ignored the contribution of 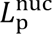 [91], implicitly setting 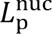 = ∞, or let 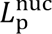 = 10 × 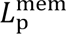 [60]. To our knowledge, our analysis is the first to provide 110 cell-nucleus paired estimates of 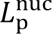 in anisotonic conditions and 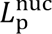 at the population level in the presence of CPAs. Our analysis shows that 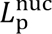 is relatively insensitive as long as 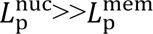 meaning that model the simplification of 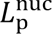 are generally appropriate for most applications. We are unaware of literature reporting nuclear *P*_s_ values and note that the presence and abundance of nuclear pores likely limit our ability to present precise estimates of nuclear CPA permeability. Nuclear pores have a diameter of 30-60 kDa allowing for facilitated permeation of small molecules like DMSO (78 Da) and EG (62 Da) [21, 80, 81]. Depending on the abundance of nuclear pores, 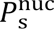 is likely high, and is likely limited in sensitivity in a similar way as nuclear hydraulic conductivity is when 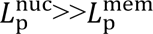. This insensitivity is reflected in the relatively high standard error (**Table 3**). Future work must consider the limiting factor the cytoplasm has on obtaining precise estimates of nuclear hydraulic conductivity.

Finally, we consider that if the cell loses ions during CPA loading, this may be beneficial as regulatory volume decrease reduces water content and therefore reducing the time required for CPA equilibration. Alternatively, it is possible that loss of cellular ion balance and possible influx of Ca2+ is detrimental to the cell leading to increased apoptosis rates. It may be more beneficial to use a media similar to cytoplasm composition. Indeed, an intracellular electrolyte composition (Euro-Collins) media with high K+ and low Ca2+ has already been shown to improve cryopreservation protocols for rat hepatocytes [92].

### 4.5. Bleb formation

Bleb formation was observed in time series images of cells in hypo-osmotic media for HepG2 monolayers (**Fig 3C**). Blebbing occurs when the actomyosin cortex is separated from the cell membrane, leading to loss of mechanical resistance[93]. Alternatively, the osmotic behavior of the cytoplasm can be explained by the leakiness of the cell membranes. In hyposmotic media, swelling causes membrane mechanosensitive channels to open in some cells, releasing macromolecules and important osmolytes [78]. Future work, however, is necessary to fully understand this phenomenon in the context of volume regulation and mass transport.

## 5. Conclusions

We investigated both equilibrium and nonequilibrium cell, cytoplasm, and nuclear volume excursions in the presence of permeating and non-permeating solutes, followed by restoration of isosmotic conditions. Using confocal microscopy, time dependent volumetric measurements were made, and were followed by image segmentation and reconstruction in 3D. This is the first measurement of its kind, using a novel approach for simultaneous measurement of cell and nuclear osmotic responses in monolayers. Our results showed that both cytoplasm and nucleus change their volumes in response to osmotic stress, which can affect the overall equilibrium volume of the cell. Thus, it is important to consider both the cell and nuclear osmotic responses when studying cell physiology for cell monolayers since both cells and nuclei are interdependent in terms of their volumetric responses and diffusion parameters. Furthermore, since nuclei deformation is associated with perturbations in biological processes, nuclear permeability estimates can provide insight into trans-nuclear mass transport but is dependent upon timescale. Altogether, a better understanding of these membrane transport characteristics will have an impact on development of more robust mathematical models as well as cryopreservation protocols tailored to the cellular state. Consequently, all these imaging methods and membrane transport parameters can be translated from adherent endothelial cells to three-dimensional tissues.

## References

1. Oswald ES, Chao PHG, Bulinski JC, Ateshian GA, Hung CT. Chondrocyte nuclear response to osmotic loading. IEEE Eng Med Biol Soc 2006. p. 3659–61.

2. Guilak F, Tedrow JR, Burgkart R. Viscoelastic properties of the cell nucleus. Biochem Biophys Res Commun. 2000;269:781–6. doi: 10.1006/BBRC.2000.2360.

3. Hu S, Chen J, Butler JP, Wang N. Prestress mediates force propagation into the nucleus. Biochem Biophys Res Commun. 2005;329:423–8. doi: 10.1016/J.BBRC.2005.02.026.

4. Wang N, Butler JP, Ingber DE. Mechanotransduction across the cell surface and through the cytoskeleton. Science. 1993;260:1124–7. doi: 10.1126/SCIENCE.7684161.

5. Ji JY, Lee RT, Vergnes L, Fong LG, Stewart CL, Reue K. Cell nuclei spin in the absence of lamin b1. J Biol Chem. 2007;282:20015–26. doi: 10.1074/JBC.M611094200.

6. Finan JD, Chalut KJ, Wax A, Guilak F. Nonlinear osmotic properties of the cell nucleus. Ann Biomed Eng. 2009;37:477. doi: 10.1007/S10439-008-9618-5.

7. Gieni RS, Hendzel MJ. Mechanotransduction from the ECM to the genome: Are the pieces now in place? J Cell Biochem. 2008;104:1964–87. doi: 10.1002/JCB.21364.

8. Nobel PS. The Boyle-Van’t Hoff relation. J Theor Biol. 1969;23:375–9. doi: 10.1016/0022-5193(69)90025-3.

9. Bush PG, Hall AC. The osmotic sensitivity of isolated and in situ bovine articular chondrocytes. Journal of Orthopaedic Research. 2001;19:768–78. doi: 10.1016/S0736-0266(01)00013-4.

10. Guilak F, Erickson GR, Ting-Beall HP. The effects of osmotic stress on the viscoelastic and physical properties of articular chondrocytes. Biophys J. 2002;82:720–7. doi: 10.1016/S0006-3495(02)75434-9.

11. Davidson AF, Glasscock C, McClanahan DR, Benson JD, Higgins AZ. Toxicity Minimized Cryoprotectant Addition and Removal Procedures for Adherent Endothelial Cells. PLoS One. 2015;10:11. doi: 10.1371/journal.pone.0142828.

12. Hempling HG, Thompson S, Dupre A. Osmotic properties of human lymphocyte. J Cell Physiol. 1977;93:293–302. doi: 10.1002/jcp.1040930215.

13. De Freitas RC, Diller KR, Lakey JRT, Rajotte R V. Osmotic behavior and transport properties of human islets in a dimethyl sulfoxide solution. Cryobiology. 1997;35:230–9. doi: 10.1006/CRYO.1997.2045.

14. Roffay C, Molinard G, Kim K, Urbanska M, Andrade V, Barbarasa V. Passive coupling of membrane tension and cell volume during active response of cells to osmosis. Proc Natl Acad Sci U S A. 2021;118:e2103228118. doi: 10.1073/PNAS.2103228118.

15. Azam I. Multiscale Transport and Osmotic Tolerance in Liver Cells and Tissues: The University of Saskatchewan; 2023.

16. Sardini A, Amey JS, Weylandt KH, Nobles M, Valverde MA, Higgins CF. Cell volume regulation and swelling-activated chloride channels. Biochim Biophys Acta Biomembr. 2003;1618:153–62. doi: 10.1016/j.bbamem.2003.10.008.

17. Corasanti JG, Gleeson D, Boyer JL. Effects of osmotic stresses on isolated rat hepatocytes. I Ionic mechanisms of cell volume regulation Am J Physiol Gastrointest Liver Physiol. 1990;258:21–2. doi: 10.1152/ajpgi.1990.258.2.g290.

18. Hoffman EK, Lambert IH, Pedersen SF. Physiology of cell volume regulation in vertebrates. Physiol Rev. 2009;89:193–277. doi: 10.1152/PHYSREV.00037.2007.

19. Okada Y. Ion channels and transporters involved in cell volume regulation and sensor mechanisms. Cell Biochem Biophys. 2004;41:233–58. doi: 10.1385/CBB:41:2:233.

20. Wehner F, Tinel H. Role of Na+ conductance, Na+-H+ exchange, and Na+-K+-2Cl-symport in the regulatory volume increase of rat hepatocytes. Journal of Physiology. 1998;506:127–42. doi: 10.1111/j.1469-7793.1998.127bx.x.

21. Paine PL, Moore LC, Horowitz SB. Nuclear envelope permeability. Nature. 1975;254:5496. doi: 10.1038/254109a0.

22. Dahl KN, Kahn SM, Wilson KL, Discher DE. The nuclear envelope lamina network has elasticity and a compressibility limit suggestive of a molecular shock absorber. J Cell Sci. 2004;117:4779–86. doi: 10.1242/JCS.01357.

23. Peters R. Nucleo-cytoplasmic flux and intracellular mobility in single hepatocytes measured by fluorescence microphotolysis. EMBO J. 1984;3:1831–6. doi: 10.1002/J.1460-2075.1984.TB02055.X.

24. Feldherr CM, Akin D. Regulation of nuclear transport in proliferating and quiescent cells. Exp Cell Res. 1993;205:179–86. doi: 10.1006/EXCR.1993.1073.

25. Finan JD, Guilak F. The effects of osmotic stress on the structure and function of the cell nucleus. J Cell Biochem. 2010;109:460. doi: 10.1002/JCB.22437.

26. Kim DH, Li B, Si F, Phillip JM, Wirtz D, Sun SX. Volume regulation and shape bifurcation in the cell nucleus. J Cell Sci. 2015;128:3375–85. doi: 10.1242/JCS.166330.

27. Richter K, Nessling M, Lichter P. Experimental evidence for the influence of molecular crowding on nuclear architecture. J Cell Sci. 2007;120:1673–80. doi: 10.1242/JCS.03440.

28. Hancock R. A role for macromolecular crowding effects in the assembly and function of compartments in the nucleus. J Struct Biol. 2004;146:281–90. doi: 10.1016/j.jsb.2003.12.008.

29. Conklin EG. Cell size and nuclear size. Journal of Experimental Zoology. 1912;12:1–98. doi: 10.1002/JEZ.1400120102.

30. Kirby TJ, Lammerding J. Emerging views of the nucleus as a cellular mechanosensor. Nat Cell Biol. 2018;20:373–81. doi: 10.1038/s41556-018-0038-y.

31. Fahy GM, MacFarlane DR, Angell CA, Meryman HT. Vitrification as an approach to cryopreservation. Cryobiology. 1984;21:407–26. doi: 10.1016/0011-2240(84)90079-8.

32. Wusteman MC, Pegg DE, Robinson MP, Wang LH, Fitch P. Vitrification media: Toxicity, permeability, and dielectric properties. Cryobiology. 2002;44:24–37. doi: 10.1016/S0011-2240(02)00002-0.

33. Benson JD, Higgins AZ, Desai K, Eroglu A. A toxicity cost function approach to optimal CPA equilibration in tissues. Cryobiology. 2018;80:144–55. doi: 10.1016/j.cryobiol.2017.09.005.

34. Model MA. Methods for cell volume measurement. Cytometry. 2018;93:281–96. doi: 10.1002/cyto.a.23152.

35. Davidson AF, Higgins AZ. Detection of volume changes in calcein-stained cells using confocal microscopy. J Fluoresc. 2013;23:393–8. doi: 10.1007/s10895-013-1202-1.

36. Fry AK, Higgins AZ. Measurement of cryoprotectant permeability in adherent endothelial cells and applications to cryopreservation. Cell Mol Bioeng. 2012;5:287–98. doi: 10.1007/s12195-012-0235-x.

37. Terry C, Dhawan A, Mitry RR, Hughes RD. Cryopreservation of isolated human hepatocytes for transplantation: State of the art. Cryobiology. 2006;53:149–59. doi: 10.1016/j.cryobiol.2006.05.004.

38. Zhou Z, Xu MJ, Gao B. Hepatocytes: a key cell type for innate immunity. Cellular & Molecular Immunology. 2016;13:3. doi: 10.1038/cmi.2015.97.

39. Sosef MN, Baust JM, Sugimachi K, Fowler A, Tompkins RG, Toner M. Cryopreservation of isolated primary rat hepatocytes: Enhanced survival and long-term hepatospecific function. Ann Surg. 2005;241:125–33. doi: 10.1097/01.sla.0000149303.48692.0f.

40. Bischof JC, Ryan CM, Tompkins RG, Yarmush ML, Toner M. Ice formation in isolated human hepatocytes and human liver tissue. ASAIO Journal. 1997;43:271–8. doi: 10.1097/00002480-199743040-00003.

41. Darr TB, Hubel A. Freezing characteristics of isolated pig and human hepatocytes. Cell Transplant. 1997;6:173–83. doi: 10.1016/S0963-6897(96)00252-7.

42. Karlsson JOM, Cravalho EG, Borel Rinkes IH, Tompkins RG, Yarmush ML, Toner M. Nucleation and growth of ice crystals inside cultured hepatocytes during freezing in the presence of dimethyl sulfoxide. Biophys J. 1993;65:2524–36. doi: 10.1016/S0006-3495(93)81319-5.

43. Massa S, Sakr MA, Seo J, Bandaru P, Arneri A, Bersini S. Bioprinted 3D vascularized tissue model for drug toxicity analysis. Biomicrofluidics. 2017. doi: 10.1063/1.4994708.

44. Sun L, Yang H, Wang Y, Zhang X, Jin B, Xie F. Application of a 3D Bioprinted Hepatocellular Carcinoma Cell Model in Antitumor Drug Research. Front Oncol. 2020. doi: 10.3389/fonc.2020.00878.

45. Zhang Y, Schattling PS, Itel F, Stadler B. Planar and cell aggregate-like assemblies consisting of microreactors and HepG2 cells. ACS Omega. 2017;2:7085–95. doi: 10.1021/acsomega.7b01234.

46. Qiu GH, Xie X, Xu F, Shi X, Wang Y, Deng L. Distinctive pharmacological differences between liver cancer cell lines HepG2 and Hep3B. Cytotechnology. 2015. doi: 10.1007/s10616-014-9761-9.

47. Duncan AW, Taylor MH, Hickey RD, Hanlon Newell AE, Lenzi ML, Olson SB. The ploidy conveyor of mature hepatocytes as a source of genetic variation. Nature. 2010;467:7316. doi: 10.1038/nature09414.

48. Toyoda H, Bregerie O, Vallet A, Nalpas B, Pivert G, Brechot C. Changes to hepatocyte ploidy and binuclearity profiles during human chronic viral hepatitis. Gut. 2005;54:297–302. doi: 10.1136/GUT.2004.043893.

49. Li H, Feng Z, Tsang TC, Tang T, Jia X, He X. Fusion of HepG2 cells with mesenchymal stem cells increases cancer-associated and malignant properties: an in vivo metastasis model. Oncol Rep. 2014;32:539–47. doi: 10.3892/OR.2014.3264.

50. Fedorchak GR, Kaminski A, Lammerding J. Cellular mechanosensing: getting to the nucleus of it all. Prog Biophys Mol Biol. 2014;115:76–92. doi: 10.1016/J.PBIOMOLBIO.2014.06.009.

51. Pachitariu M, Stringer C. Cellpose 2.0: how to train your own model. Nature Methods. 2022;19:12. doi: 10.1038/s41592-022-01663-4.

52. Stringer C, Wang T, Michaelos M, Pachitariu M. Cellpose: a generalist algorithm for cellular segmentation. Nature Methods. 2020;18:1. doi: 10.1038/s41592-020-01018-x.

53. Mazur P. Life in the Frozen State. Boca Raton: CRC Press; 2004.

54. Benson JD. Some comments on recent discussion of the Boyle van’t Hoff relationship. Cryobiology. 2012;64:118–20. doi: 10.1016/j.cryobiol.2011.12.001.

55. Prickett RC, Elliott JAW, Hakda S, McGann LE. A non-ideal replacement for the Boyle van’t Hoff equation. Cryobiology. 2008;57:130–6. doi: 10.1016/J.CRYOBIOL.2008.07.002.

56. Jacobs MH, Stewart DR. A simple method for the quantitative measurement of cell permeability. J Cell Comp Physiol. 1932;1:71–82. doi: 10.1002/JCP.1030010107.

57. Benson JD, Benson CT, Critser JK. Mathematical model formulation and validation of water and solute transport in whole hamster pancreatic islets. Math Biosci. 2014;254:64–75. doi: 10.1016/J.MBS.2014.06.003.

58. Benson JD. Mathematical modeling and optimization of cryopreservation in single cells. Methods in Molecular Biology: Humana Press Inc.; 2021.

59. Abazari A, Elliott JAW, Law GK, McGann LE, Jomha NM. A biomechanical triphasic approach to the transport of nondilute solutions in articular cartilage. Biophys J. 2009;97:3054–64. doi: 10.1016/j.bpj.2009.08.058.

60. Hoffmann PC, Kim H, Obarska-Kosinska A, Kreysing JP, Andino-Frydman E, Cruz-León S. Nuclear pore permeability and fluid flow are modulated by its dilation state. Mol Cell. 2025;85:537–54.e11. doi: 10.1016/j.molcel.2024.11.038.

61. Wehner F, Lawonn P, Tinel H. Ionic mechanisms of regulatory volume increase (RVI) in the human hepatoma cell-line HepG2. Pflugers Arch. 2002;443:779–90. doi: 10.1007/s00424-001-0765-x.

62. Zelenina M, Brismar H. Osmotic water permeability measurements using confocal laser scanning microscopy. Eur Biophys J. 2000;29:165–71. doi: 10.1007/pl00006645.

63. Gregg JL, McGuire KM, Focht DC, Model MA. Measurement of the thickness and volume of adherent cells using transmission-through-dye microscopy. Pflugers Arch. 2010;460:1097–104. doi: 10.1007/S00424-010-0869-2.

64. Peckys DB, Kleinhans FW, Mazur P. Rectification of the Water Permeability in COS-7 Cells at 22, 10 and 0°C. PLoS One. 2011;6:e23643. doi: 10.1371/JOURNAL.PONE.0023643.

65. Olver DJ, Azam I, Benson JD. HepG2 cells undergo regulatory volume decrease by mechanically induced efflux of water and solutes. Biomech Model Mechanobiol. 2024;23:1781–99. doi: 10.1007/s10237-024-01868-w.

66. Li L, Chen Z, Zhang M, Panhwar F, Gao C, Zhao G. Cell membrane permeability coefficients determined by single-step osmotic shift are not applicable for optimization of multi-step addition of cryoprotective agents: As revealed by HepG2 cells. Cryobiology. 2017;79:82–6. doi: 10.1016/j.cryobiol.2017.09.006.

67. Azam I, Benson JD. Silymarin mediated osmotic responses and damage in HepG2 cell suspensions and monolayers. Cryobiology. 2023;112:104552. doi: 10.1016/j.cryobiol.2023.10455.

68. Jiang H, Sun SX. Cellular pressure and volume regulation and implications for cell mechanics. Biophys J. 2013;105(3):609–19. doi: 10.1016/j.bpj.2013.06.021. PubMed PMID: 23931309; PubMed Central PMCID: PMCPMC3736675.

69. Hoffmann EK. The pump and leak steady-state concept with a variety of regulated leak pathways. J Membr Biol. 2001;184(3):321–30. Epub 20020205. doi: 10.1007/s00232-001-0102-5. PubMed PMID: 11891558.

70. Hoffmann EK, Lambert IH, Pedersen SF. Physiology of cell volume regulation in vertebrates. Physiol Rev. 2009;89(1):193–277. doi: 10.1152/physrev.00037.2007. PubMed PMID: 19126758.

71. Keener J, Sneyd J. Mathematical Physiology. New York: Springer-Verlag; 1998.

72. Cruzeiro-Hansson L, Mouritsen OG. Passive ion permeability of lipid membranes modelled via lipid-domain interfacial area. Biochim Biophys Acta. 1988;944(1):63–72. doi: 10.1016/0005-2736(88)90316-1. PubMed PMID: 3415999.

73. Quinn PJ. A lipid-phase separation model of low-temperature damage to biological membranes. Cryobiology. 1985;22(2):128–46. doi: 10.1016/0011-2240(85)90167-1. PubMed PMID: 3920005.

74. Blok MC, van der Neut-Kok EC, van Deenen LL, de Gier J. The effect of chain length and lipid phase transitions on the selective permeability properties of liposomes. Biochim Biophys Acta. 1975;406(2):187–96. doi: 10.1016/0005-2736(75)90003-6. PubMed PMID: 1191647.

75. He F, Liu W, Zheng S, Zhou L, Ye B, Qi Z. Ion transport through dimethyl sulfoxide (DMSO) induced transient water pores in cell membranes. Mol Membr Biol. 2012;29(3-4):107–13. Epub 20120601. doi: 10.3109/09687688.2012.687460. PubMed PMID: 22656651.

76. de Menorval MA, Mir LM, Fernandez ML, Reigada R. Effects of dimethyl sulfoxide in cholesterol-containing lipid membranes: a comparative study of experiments in silico and with cells. PLoS One. 2012;7(7):e41733. Epub 20120725. doi: 10.1371/journal.pone.0041733. PubMed PMID: 22848583; PubMed Central PMCID: PMCPMC3404987.

77. Fernandez ML, Reigada R. Effects of dimethyl sulfoxide on lipid membrane electroporation. J Phys Chem B. 2014;118(31):9306–12. Epub 20140725. doi: 10.1021/jp503502s. PubMed PMID: 25035931.

78. Casula E, Asuni GP, Sogos V, Fadda S, Delogu F, Cincotti A. Osmotic behaviour of human mesenchymal stem cells: Implications for cryopreservation. PLoS One. 2017. doi: 10.1371/JOURNAL.PONE.0184180.

79. Christmann J, Azer L, Dörr D, Fuhr GR, Bastiaens P, Wehner F. Adaptive responses of cell hydration to a low temperature arrest. J Physiol. 2016;594:1663–76. doi: 10.1113/JP271245.

80. Timney BL, Raveh B, Mironska R, Trivedi JM, Kim SJ, Russel D. Simple rules for passive diffusion through the nuclear pore complex. J Cell Biol. 2016. doi: 10.1083/JCB.201601004.

81. Samudram A, Mangalassery BM, Kowshik M, Patincharath N, Varier GK. Passive permeability and effective pore size of HeLa cell nuclear membranes. Cell Biol Int. 2016;40:991–8. doi: 10.1002/CBIN.10640.

82. Wang R, Brattain MG. The maximal size of protein to diffuse through the nuclear pore is larger than 60kDa. FEBS Lett. 2007;581:3164. doi: 10.1016/J.FEBSLET.2007.05.082.

83. Popken P, Ghavami A, Onck PR, Poolman B, Veenhoff LM. Size-dependent leak of soluble and membrane proteins through the yeast nuclear pore complex. Mol Biol Cell. 2015;26:1386–94. doi: 10.1091/MBC.E14-07-1175.

84. Alberts B, Johnson A, Lewis J, Raff M, Roberts K, Walter P. The Transport of Molecules between the Nucleus and the Cytosol. Molecular Biology of the Cell. New York: Garland Science; 2002.

85. Wu Y, Pegoraro AF, Weitz DA, Janmey P, Sun SX. The correlation between cell and nucleus size is explained by an eukaryotic cell growth model. PLoS Comput Biol. 2022. doi: 10.1371/JOURNAL.PCBI.1009400.

86. Cantwell H, Nurse P. Unravelling nuclear size control. Current Genetics. 2019;65:6. doi: 10.1007/S00294-019-00999-3.

87. Ganguly A, Bhattacharjee C, Bhave M, Kailaje V, Jain BK, Sengupta I. Perturbation of nucleo-cytoplasmic transport affects size of nucleus and nucleolus in human cells. FEBS Lett. 2016;590:631–43. doi: 10.1002/1873-3468.12077.

88. Kume K, Cantwell H, Neumann FR, Jones AW, Snijders AP, Nurse P. A systematic genomic screen implicates nucleocytoplasmic transport and membrane growth in nuclear size control. PLoS Genet. 2017. doi: 10.1371/JOURNAL.PGEN.1006767.

89. Albro MB, Chahine NO, Caligaris M, Wei VI, Likhitpanichkul M, Ng KW. Osmotic Loading of Spherical Gels: A Biomimetic Study of Hindered Transport in the Cell Protoplasm. J Biomech Eng. 2007;129:503. doi: 10.1115/1.2746371.

90. Jaiswal A, Hoerth CH, Zúñiga Pereira AM, Lorenz H. Improved spatial resolution by induced live cell and organelle swelling in hypotonic solutions. Sci Rep. 2019;9:12911. doi: 10.1038/s41598-019-49408-2.

91. Horowitz SB. The permeabaility of the amphibian oocyte nucleus, in situ. J Cell Biol. 1972;54:609–25. doi: 10.1083/jcb.54.3.609.

92. De Loecker R, Fuller BJ, Gruwez J, De Loecker W. The effects of cryopreservation on membrane integrity, membrane transport, and protein synthesis in rat hepatocytes. Cryobiology. 1990;27:143–52. doi: 10.1016/0011-2240(90)90006-p.

93. Norman LL, Bruges J, Sengupta K, Sens P, Aranda-Espinoza H. Cell blebbing and membrane area homeostasis in spreading and retracting cells. Biophys J. 2010;99:1726–33. doi: 10.1016/J.BPJ.2010.07.031.

